# Evaluating the relationships between the legal and illegal international wildlife trades

**DOI:** 10.1101/726075

**Authors:** Derek P. Tittensor, Michael Harfoot, Claire McLardy, Gregory L. Britten, Katalin Kecse-Nagy, Bryan Landry, Willow Outhwaite, Becky Price, Pablo Sinovas, Julian Blanc, Neil D. Burgess, Kelly Malsch

## Abstract

The international legal trade in wildlife can provide economic and other benefits, but when unsustainable can be a driver of population declines. This impact is enhanced by the additional burden of illegal trade. We combined law-enforcement time-series of seizures of wildlife goods imported into the United States (US) and the European Union (EU) with data on reported legal trade to evaluate the evidence for any relationships. Our analysis examined 28 (US) and 20 (EU) taxon-products with high volume and frequency of reported trade and seizures. On average, seizures added 28% and 9% to US and EU reported legal trade levels respectively, and in several cases exceeded legal imports. We detected a significant but weak overall positive relationship between seizure volumes and reported trade imported to the US over time, but none in the EU. Our findings suggest a complex and nuanced temporal association between the illegal and legal wildlife trades.

## Introduction

The international trade in wildlife is a long-standing activity (Jenkins & Broad 1994) with the potential for substantial economic benefits. However, with the recognition that unsustainable trade can be damaging to source animal and plant populations, the Convention on International Trade in Endangered Species of Wild Fauna and Flora (CITES) entered into force in 1975 as a global policy mechanism to balance benefits and impacts and ensure sustainability. Over 35,000 species are listed in CITES, in one of three Appendices with differing trade implications. The collation of CITES permits into the CITES Trade Database (trade.cites.org) has enabled an overview of the legally regulated and reported wildlife trade at global (Harfoot et al. 2018) and regional (Sinovas et al. 2016, 2017; Outhwaite & Brown 2018) scales, as well as specific studies on individual taxa (Wu 2016).

Knowledge of the illegal wildlife trade remains much more limited due to its covert nature and the necessity for detection and interdiction. The UN Office on Drugs and Crime identified distinct markets for the trafficking of specific wildlife products and product groups (UNODC 2016), and facets of illegal trade have been examined through conducting in-depth studies (Beastall et al. 2016). However, the broader relationship between the relative volumes of legal and illegal wildlife trade remains ill-defined. Analyses of seizures across multiple taxa remain rare (but see Mundy-Taylor 2013), as do comparisons to legal trade flows.

Here we analyse seizures of CITES-listed taxon-products by law-enforcement agencies in the United States (US) and the European Union (EU), primarily representing interdiction at borders, and evaluate their association with reported legal trade volumes. A ‘taxon-product’ represents a taxon (typically species) and product(s) that are traded equivalently in a destination market (for example, leather goods from *Caiman crocodilus* and its subspecies). As trends in seizures may reflect variation in enforcement as well as illegal trade (Underwood et al. 2013), we included co-variates representing enforcement effort. We focussed on taxon-products with high volumes and frequency of reported trade and seizures. Seizure volumes represent an ‘on-the-ground’ measure of contraband goods interceptions.

Our study had three main objectives. Firstly, to compare relative volumes of reported legal trade and seizures for products entering the US and the EU. This provides a ‘minimum estimate’ of illegal trade and how it compares to the legal trade in these jurisdictions. Secondly, to test for statistically significant associations over time, either positive or negative, between reported legal trade and seizure volumes. Thirdly, to assess the utility of these data sources to help inform species conservation and policy responses.

## Methods

### Data sources and extraction

Importer-reported wildlife trade data from three sources were combined for this analysis: the CITES Trade Database (trade.cites.org) for the reported legal trade, and the Law Enforcement Management Information System (LEMIS) and EU Trade in Wildlife Information Exchange (EU-TWIX) databases for US and EU seizures respectively. Seizures primarily reflect interdictions at international borders.

All records of importer-reported legal trade in CITES-listed taxa to the US and the EU for 1975 – 2014 were extracted from the CITES Trade Database, with code “I” (“confiscated or seized”) records excluded. The final year (2014) was selected to ensure high record completeness across all databases. We analysed records from all sources of trade and for all purposes, aggregated across exporting countries. To ensure consistency with EU-TWIX, EU records were filtered to exclude those involving trade between Member States (i.e. analysis was of trade entering the EU from outside its borders).

For US seizures, we filtered shipment records to those where the importing Party was the US and the source was coded as “confiscated or seized” (“I”). Data were extracted for the same time-period as the reported trade, though most time-series began later, with LEMIS becoming operational in 1983. For EU seizures, we analysed the period 2005 to 2014; this represented the longest period over which relatively complete seizure information was available from the EU-TWIX database. Data were standardised according to CITES standard nomenclature.

Records were aggregated to unique taxon (species or sub-species), product and unit combinations (‘taxon-products’) to yield time-series of reported legal trade volumes and seizures. As our focus was on testing for consistency of relationships between reported trade and seizures, we analysed taxon-products which were frequently interdicted, had frequent legal trade across multiple years, and for which trade volumes and seizures were relatively high; see Appendix S1 for full criteria and details.

We removed products for which the number or amount of wildlife they contained could not be ascertained (medicines, powder, waxes, and derivatives). Most plants were traded as medicines or other derivatives, and/or only identified to family level, hence were omitted from the analysis. Wildlife subspecies and product types that are traded equivalently were combined (see Appendix S2 and Table S1). Species transferred between CITES Appendices during the analysis period were excluded, to prevent confounding effects on the association between reported trade volumes and seizures. Taxa that were split-listed (populations on both Appendix I and II) and sub-species listed in a different Appendix to their parent were similarly excluded; thus some iconic species such as African elephant (*Loxodonta africana*) were not included in this analysis. One exception to the above was caviar (from *Acipenseriformes*), which was retained given the importance of the product and its interchangeability.

Following quality control processes, we were left with 28 time-series for the US and 20 for the EU, representing combined taxon-products which were equivalent among databases, identified to species or genus level, with relatively high volumes and frequencies of both trade and seizures. We then modelled the relationship between reported legal trade and seizures over time for each taxon-product and destination market.

### Modelling approach for relating reported trade to law enforcement seizures

We used a hierarchical Bayesian modelling approach to assess the relationship between seizure and reported legal trade volumes, with a unique slope parameter for each taxon-product. We considered a slope (representing the change in illegal seizures with legal trade) as statistically significant if its 95% posterior probability interval did not contain zero. A significant positive slope implied that reported legal trade increased with seizures volumes also increased, and vice versa. If the slope estimate was while a significantly negative slope implied the opposite Finally, if the slope estimate was not significantly different from zero, there was no evidence for a relationship in the destination market; this was our null expectation. We compared these slope values across taxon-products, markets, CITES Appendices, and taxonomic groups.

To account for variation in enforcement effort, we included the total number of seizures across all taxon-products in each year in the US or the EU as a co-variate. While seizures of specific individual taxon-products may fluctuate, the total number of seizures is more likely to reflect enforcement effort than total illegal volume. Thus, we assume that this proxy reflects on the ground (border) enforcement effort reasonably well. We checked this assumption in the US through comparing results to an alternative (but shorter) proxy of the total number of containers inspected each year; however, we did not use this as the primary effort proxy as the time-series was much shorter and not available for the EU. We also accounted for differences in seizure reporting rates to CITES between EU Member States with an additional covariate (Table S2).

All models were fitted within a Bayesian modelling framework using a Hamiltonian Monte Carlo approach (Stan Development Team 2015), and tested on simulated data prior to being fitted to time-series to ensure that they could recapture known parameters. See the Supplementary Methods for full details.

## Results

Figure 1 shows example time-series for importer-reported legal trade and law-enforcement seizure volumes from the US and the EU. All time-series are displayed in Figures S1 and S2.

**Figure 1:**
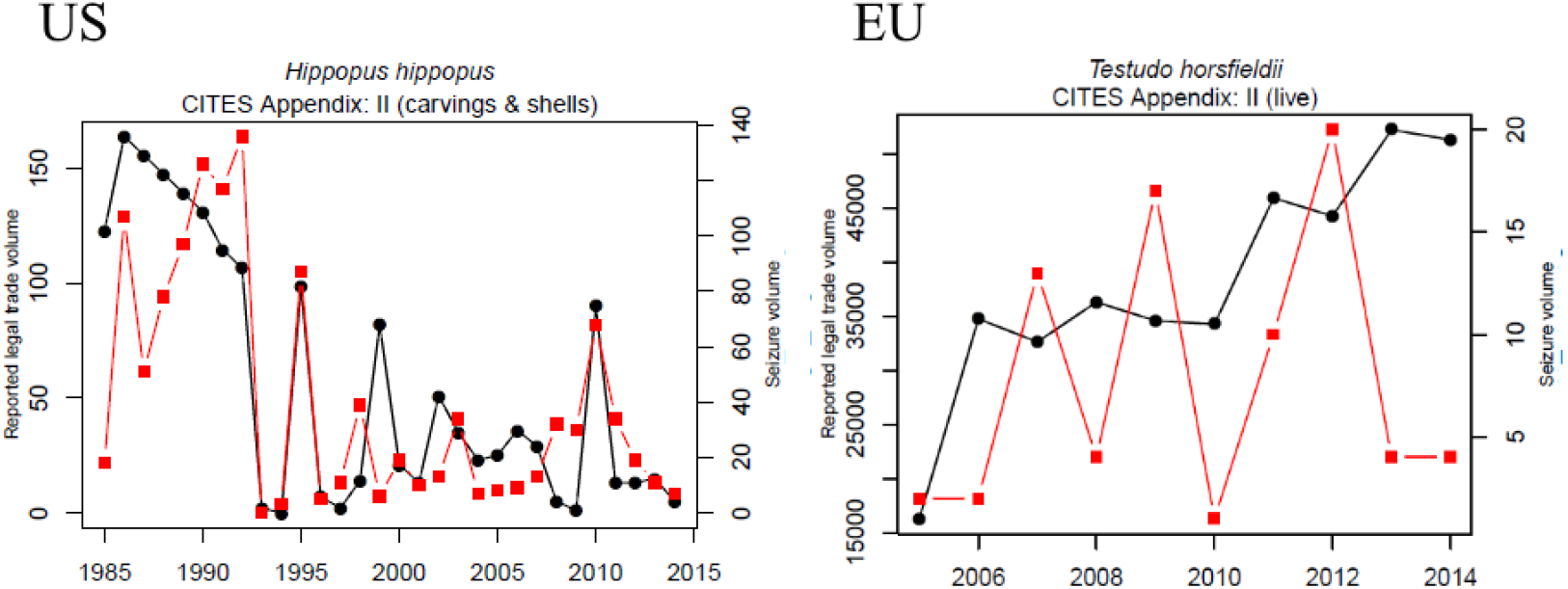
Example time-series plots for US and EU CITES reported legal trade and law-enforcement seizure volumes of wildlife products. Black circles and lines represent reported trade volume, and red squares and lines seizure volumes. Data have been processed using a robust statistics approach as described in the Methods. CITES data are importer-reported and include trade from all sources except source “I” (confiscated or seized). For full time-series and model fits see Figures S1 and S2.

### Relationship between seizure volumes and reported legal trade volumes

The ratios of reported legal trade volumes to seizure volumes summed over the period of analysis are shown in Figures 2 (US) and 3 (EU). There was a wide range of variability across taxon-products, from seizures adding almost no additional volume, to being equivalent to or even exceeding reported legally-traded items. In the US, the mean ratio of seizures to reported trade was 0.28 to 1 (i.e. seizures added, on average, around 28% to the volume of reported legal trade). CITES Appendix-I listed species generally had much higher ratios, potentially because legal trade is only permitted in exceptional circumstances (e.g. for scientific purposes). In the EU, the mean ratio was around 0.09 to 1 (i.e. seizures added almost 10% to the total reported trade). However, for some EU taxon-products it was substantially higher: for instance, seizure volumes of Appendix-II listed *Tridacna maxima* shells during this period was close to the total reported legal trade volume (Fig. 3).

**Figure 2:**
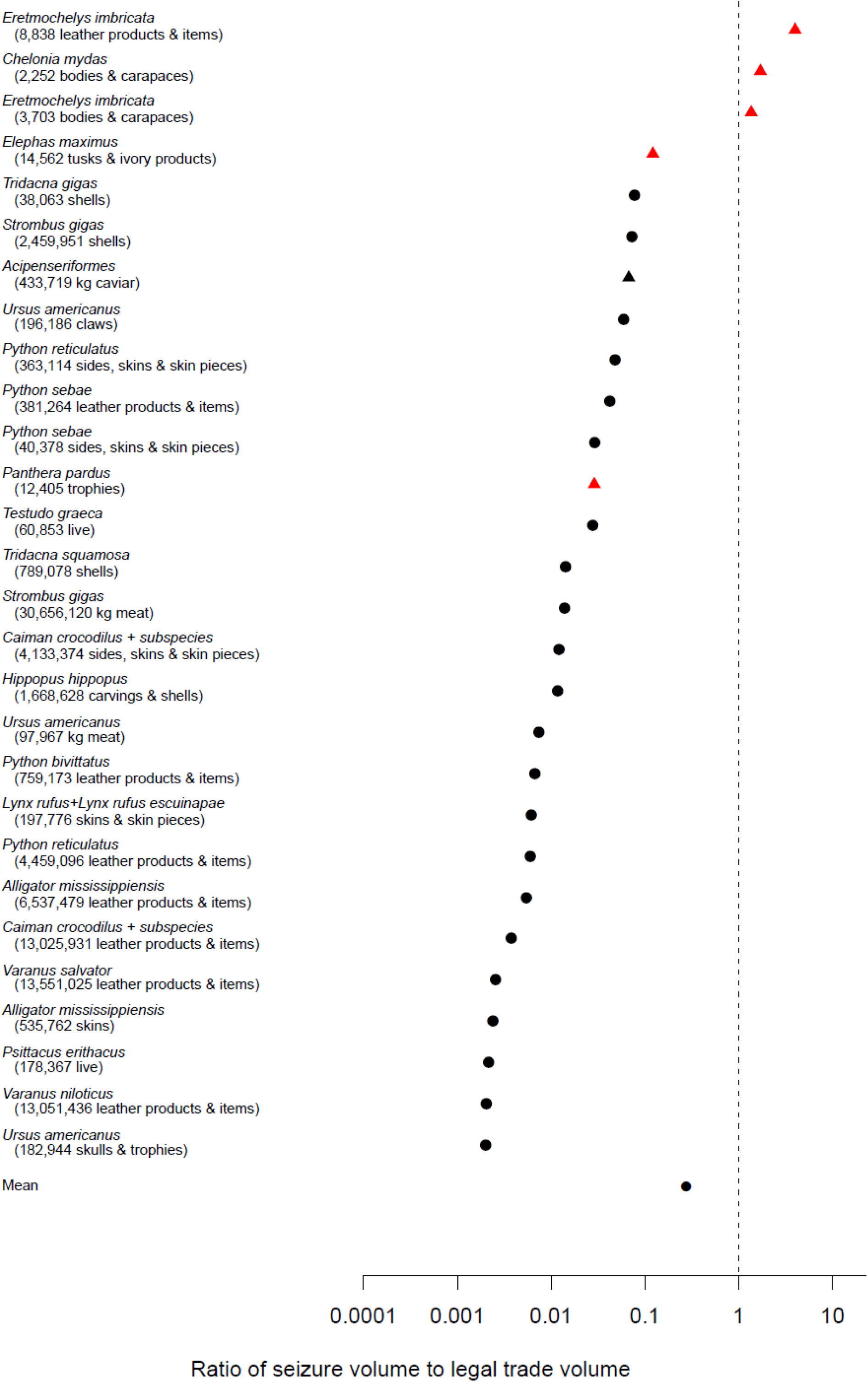
Relative ratios of seizures to importer-reported legal CITES trade (from all sources) for the US, summed over the period of analysis (typically 1980s to 2014). Symbols indicate the ratio of seizures to reported trade volume (in terms of numbers or kilograms of products). Vertical dotted line indicates a 1:1 ratio at which the number or weight of reported seized goods would be equivalent to the total reported trade reported during this period. Numbers in brackets indicate the summed number/weight of both reported and seized goods for each taxon-product. Red triangles indicate CITES Appendix I taxa; black circles indicate Appendix II taxa. Note that Acipenseriformes includes one Appendix I and two Appendix II species. Taxon-products ordered by relative ratio, from largest (top) to smallest (bottom).

**Figure 3:**
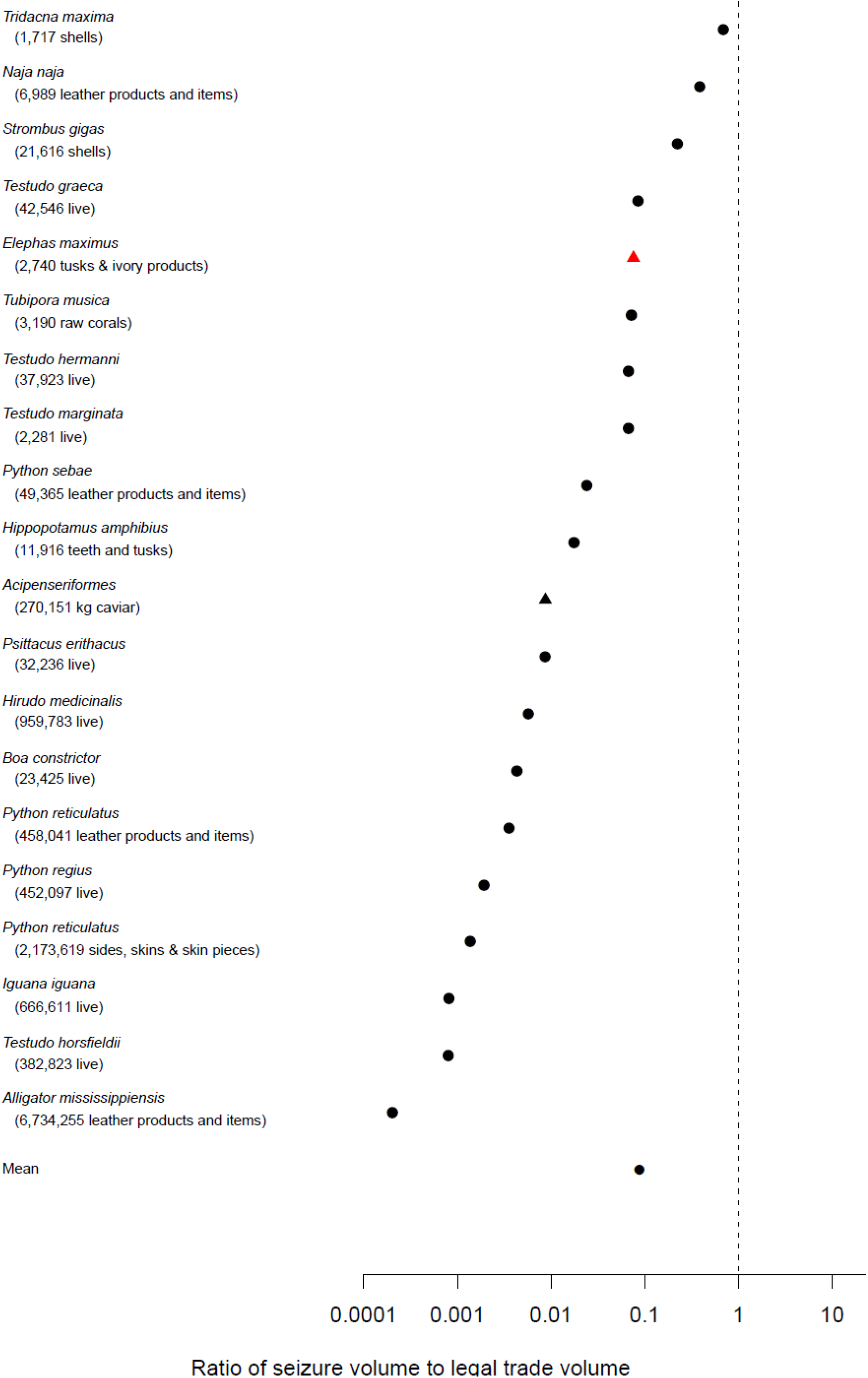
Relative ratios of seizures to importer-reported legal CITES trade (from all sources) for the EU, summed over the period of analysis (2005 to 2014). Symbols indicate the ratio of seizures to reported trade volume (in terms of numbers or kilograms of products). Vertical dotted line indicates a 1:1 ratio at which the number or weight of reported seized goods would be equivalent to the total reported trade reported during this period. Numbers in brackets indicate the summed number/weight of both reported and seized goods for each taxon-product. Red triangles indicate CITES Appendix I taxa; black circles indicate Appendix II taxa. Note that Acipenseriformes (black triangle) includes one Appendix I and two Appendix II species. Taxon-products ordered by relative ratio, from largest (top) to smallest (bottom).

### Association between trends in legal trade and seizures over time

In the US, the meta-analytic median indicated an overall significant and positive relationship between reported legal trade and seizure volumes across all taxon-products included in the analysis (Fig. 4). Furthermore, 5 of 28 (18%) individual taxon-product combinations showed a significant positive relationship between reported trade and seizures. For the EU, none of the 20 taxon-product combinations showed a significant relationship and the meta-analytic median was similarly non-significant (Fig. 5). The variance accounted for was in general fairly low (Figs. S1 and S2) for taxon-products in both destination markets. There were no significant differences in meta-analytic posterior slope estimates between taxonomic groups, product types, and CITES Appendices.

**Figure 4:**
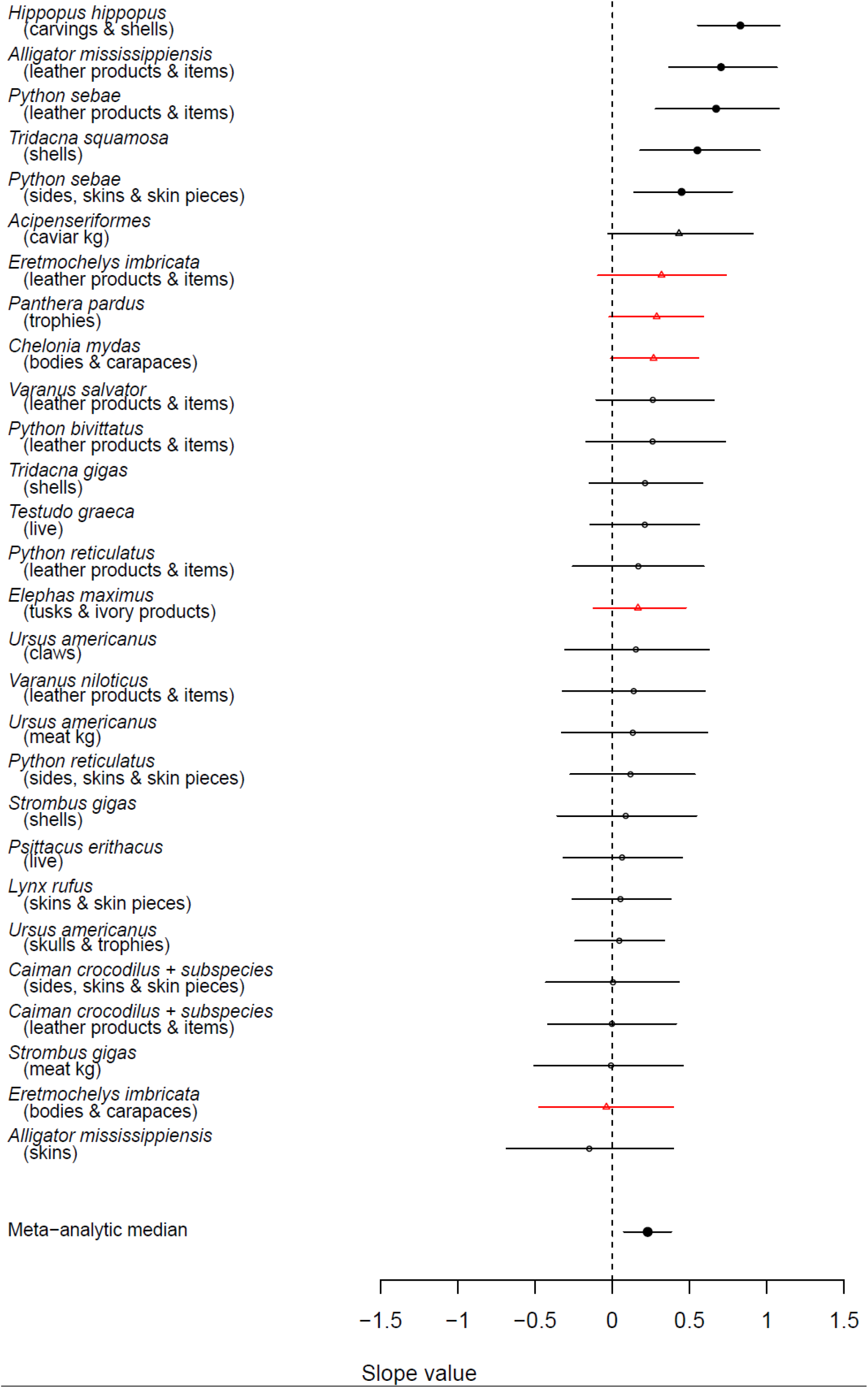
Strength of the relationship between the importer-reported legal trade volume from all sources and seizure volume over time for taxon-products in the US (time-series typically early 1980s to 2014). Values are the slope parameter from a hierarchical Bayesian regression model relating reported trade as a predictor of seizure volume (with units increase in seizures per unit change in legal trade). A positive value indicates a positive relationship between reported trade and seizure volumes, while a negative value indicates an inverse relationship. The meta-analytic median represents the hierarchical slope parameter value. Symbols indicate posterior Bayesian median model estimates, while lines indicate 95% credible intervals. Filled symbols indicate a value that is significantly different from zero while open circles indicate a non-significant relationship. Red triangles indicate CITES Appendix I species; all other species CITES Appendix II during the analysis period. Note that Acipenseriformes (black triangle) includes one Appendix I and two Appendix II species. Taxa ordered by effect size.

**Figure 5:**
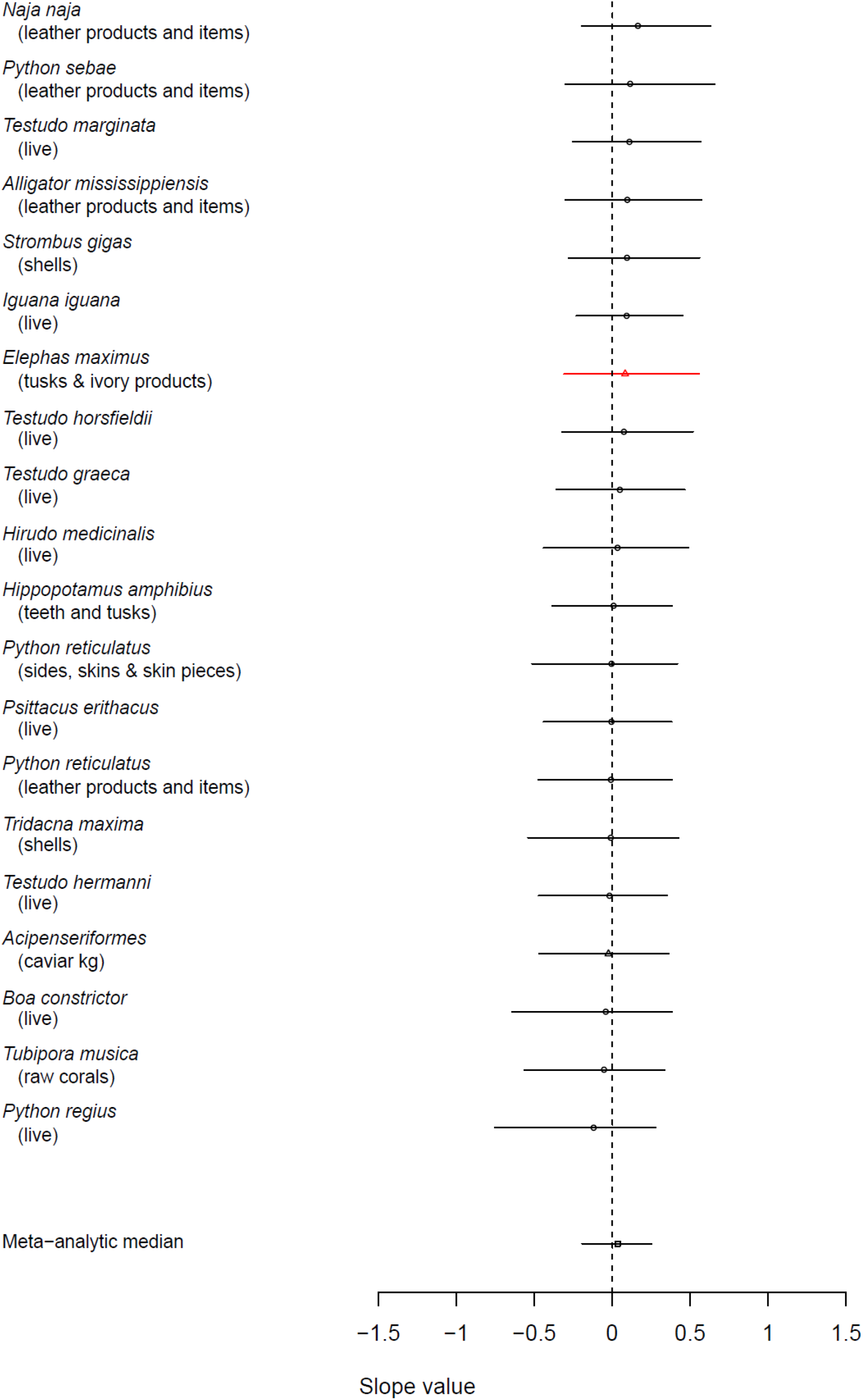
Strength of the relationship between the importer-reported legal trade volume from all sources and seizure volume over time for taxon-products in the EU (2005 to 2014). Values are the slope parameter from a hierarchical Bayesian regression model relating reported trade as a predictor of seizure volume (with units increase in seizures per unit change in legal trade). A positive value indicates a positive relationship between reported trade and seizure volumes, while a negative value indicates an inverse relationship. The meta-analytic median represents the hierarchical slope parameter value. Symbols indicate posterior Bayesian median model estimates, while lines indicate 95% credible intervals. Filled symbols indicate a value that is significantly different from zero while open circles indicate a non-significant relationship. Red triangles indicate CITES Appendix I species; all other species CITES Appendix II during the analysis period. Note that Acipenseriformes (black triangle) includes one Appendix I and two Appendix II species. Taxa ordered by effect size.

Testing robustness by using an alternative proxy for enforcement effort in the US (Fig. S3) suggested broadly similar patterns, though while the meta-analytic median was still positive it was non-significant, and individual time-series showed differences. This may be due to the shorter time-series available for the alternative effort proxy, as shortening the original effort time-series to the same length also resulted in a non-significant outcome (Fig. S4).

## Discussion

### Relationship between seizure volumes and legal trade volumes

Seizures represent an absolute lower bound on the level of illegal trade activity. Even so, the relative volume of seizures to reported legal trade was substantial (0.28 to 1 for the US and 0.09 to 1 for the EU). Differences between US and EU destination markets may be due to longer US time-series being, differences in taxon-products analysed, or variation in enforcement targeting and effort.

Our analysis highlights multiple taxa (Figs. 2 and 3) for which the relative volume of seized products was very high, and in some cases higher than the legally reported trade. The export of Appendix I and II taxa requires a “non-detriment finding” (NDF) to be made to ensure that trade is sustainable and not detrimental to the survival of the species in the wild. Scientific Authorities should ‘consider the volume of legal and illegal trade’ as part of their NDF (CITES Resolution Conf. 16.7 Rev. CoP17). Assessing relative volumes of reported legal trade and seizures can help to identify and prioritize species where the illegal trade may constitute a significant proportion of all trade. However, unless there is a single population of a taxon involved, it may be challenging to translate from the seizures (potentially from multiple populations/countries) to the harvest of individual populations at the national level where NDFs are made. Furthermore, different parts of the same specimen can enter the illegal trade as separate products (such as meat, claws, skulls/trophies), meaning that it is not as straightforward as simply adding up the number of seized specimens to assess impact on wild populations.

### Association between trends in legal trade and seizure volumes over time

The analysis of relationships between reported and seized wildlife goods suggests a complex and nuanced picture (Figs. 4 and 5). The US data showed a significant overall positive relationship, as did almost 20% of individual taxon-products. However, while always positive, the relationship was only significant with the full longer-term effort proxy (Figs. S3 and S4). There were no significant positive associations for the EU, and no significant negative associations in either market. We note that the EU time-series (10 years) were short compared to the US time-series (up to ~30 years), and shorter time-series may be noisier and statistically less likely to detect long-term trends.

However, in general it appears that there may be a weakly positive relationship (though not necessarily causality) between reported trade and seizures for the US, but with high variability among taxon-products. This could be due to multiple factors. An increase in reported legal trade volume for a specific taxon-product could lead to greater awareness among authorities, and increased enforcement effort or ability to identify illegal specimens. Administrative errors that result in seizures may also increase as legal volume increases. Some taxon-products are traded as personal items and may have been transported without necessary paperwork due to a lack of awareness rather than as part of trafficking operations. Increased legal trade may also lead to an increase in illegal trade if demand cannot be met (although conversely, increased reported trade could also provide an adequate supply to the market thus reducing the need for illegally sourced products). Finally, there may be non-casual but correlated factors; for example, more direct flights between the United States and source countries over the multi-decadal span of the analysis may have increased the potential for both legal and illegal trade.

### Policy, management, and conservation implications

The policy and wildlife management/conservation implications of this study are three-fold. Firstly, the volume of additional trade that seizures represent over and above reported trade (Figs. 2 and 3) implies that these data, notwithstanding the scaling difficulties outlined above, should be considered when estimating impacts of trade and deciding permissible trade levels. We have identified specific taxa for which seizures data imply a much greater volume of trade than apparent from the reported legal trade alone, suggesting impacts on these taxa may be underestimated. Ensuring that timely information on seizures is communicated to source countries may assist in the determination of NDFs and assessment of impact of overall trade levels.

Secondly, there were specific taxon-products for which significant positive relationships between reported legal trade volume and seizure volume were detected (Fig. 4), as well as an overall significant but weakly positive relationship across all taxon-products in the US. Further exploration is warranted as to why this might be the case. Additional variables that may influence outcomes (e.g. countries involved in trade, value and detectability of commodities) may help to shed light on this. Careful analysis of specific taxon-products (UNODC 2016) may be helpful in this respect, though only meta-analyses can detect broader general trends across taxa.

The third policy implication is that collecting and centralizing data on seizures, enforcement effort, reported legal trade, and reporting effort is essential for understanding the relationship between the legal and illegal wildlife trades. Given this, we anticipate that the value of both the US and EU datasets to infer patterns will only grow over time. Since 2017, CITES Parties have been urged to report on illegal trade in accordance with Resolution Conf. 11.17 (Rev. CoP17), which may provide additional understanding of illegal wildlife trade flows. In addition, identifying and centralising effective enforcement effort proxies, maintaining and extending current time-series, and locating such data for more target markets will all help to build a more complete picture of the global wildlife trade.

### Assumptions and caveats

The existence of a statistical association between reported legal trade volume and seizure volume is not necessarily indicative of a causal relationship between the legal and illegal wildlife trades. Variation in seizures of specific taxon-products over time can be due to many factors (including unmeasured), and although we have adjusted for biases in enforcement and reporting effort, the proxies used for this purpose are imperfect.

An important aspect of any observed relationship is that an evolving policy and interdiction environment can impact enforcement effort and targeting over time. For example, EU import restrictions for birds since 2005 could potentially cause a change in dynamics (though our EU analysis only included data starting in 2005). The EU also has a system of import restrictions for specific species-country combinations. For example, imports of wild *Testudo horsfieldi* from Tajikistan were suspended in 2005, but the suspension was removed in 2008. However, in this analysis we examined the ‘average’ trend over time (across all exporters) and attempted to minimize the impacts of policy decisions and changes in listing status by only analysing taxon-products which remained in the same CITES Appendix and EU Annex.

Finally, we note that our analysis only extends to taxon-products with relatively high seizure and reported trade volumes, and identification down to species level in all databases. Patterns and relationships may vary for other taxa, products, or destination markets.

## Conclusions

Our analysis of trade into the US and EU identified taxon-products for which seizures represent a substantial, and in some cases a majority, of trade volume. This highlights the importance of accounting for known illegal trade (i.e. seizures), where feasible for individual populations, as a ‘lower bound’ on overall trade volume when making non-detriment findings.

In the US, there was a significant positive overall relationship between reported legal trade and seizures over time, though highly variable among taxon products and contingent on effort proxy used. We did not detect a general pattern in the EU. Our findings suggest that for some markets and taxon-products there may be a weakly positive association between legal trade and seizure volumes, though it remains complex and merits further evaluation of robustness. We have also identified the critical importance of standardised long-term monitoring data, including on enforcement effort, to enable assessments of the legal and illegal wildlife flows around the globe.

Our analysis reinforces the complexities of the illegal wildlife trade and provides a starting point to further explore the nuances of specific taxa, products, enforcement environments, and destination markets. Illegal wildlife trade is a global challenge that can have profoundly deleterious consequences for targeted populations but remains enormously difficult to quantify. Long-term datasets and analyses on border seizures of wildlife products and their relationship to reported trade may help to shed light on the scale, trends, and threat of this largely covert activity.

## Acknowledgements and data

The authors acknowledge UN Environment in supporting this analysis, and the EU Member States for allowing us to analyse the seizures data extracted from the EU-TWIX database. GLB acknowledges funding from CBIOMES Grant # 549931.We thank the CITES Secretariat, and Ted Leggett from the UN Office on Drugs and Crime for helpful inputs, Vinciane Sacré and Magdalena Norwisz for providing data and information, Thomasina Oldfield for reviewing the manuscript, and Fiona Underwood, Bob Carpenter, and Mitzi Morris for help and advice on the statistical framework.

