## Supplementary Material for "Evaluating the relationships between the legal and illegal international wildlife trades"

### Appendix S1: Full Methods

*Data sources, extraction, and aggregation*

Data from three sources were combined for this analysis: the CITES Trade Database (trade.cites.org) for the reported (assumed legal) trade, and the Law Enforcement Management Information System (LEMIS) and EU Trade in Wildlife Information Exchange (EU-TWIX) databases for US and the EU seizures respectively. Seizure records within the LEMIS Database consist primarily of federal border seizures, whilst for EU-TWIX 87% of seizures were made at EU borders (i.e. not between two EU Member States or internally). We note here that seizures therefore primarily reflect interdictions of wildlife products at international or supra-national borders, whereas other types of illegal trade (such as mis-declaring specimens as captive-bred when wild trade is illegal) will not be captured by these data.

The CITES Trade Database contains all reported international trade in CITES-listed taxa exported from or imported by CITES Parties as described in their official annual reports to the Convention. Since seizure data from LEMIS and EU-TWIX are predominantly reported by importing counties, we used importer reported CITES records for consistency. All records of trade in CITES-listed taxa for the years 1975 – 2014 at the time of analysis were extracted (~16.75 million records) and all legal trade retained (i.e. code "I" "confiscated or seized" records excluded). For consistency with seizures, we analysed records from all sources of trade (e.g. captive or wild-sourced), for all purposes (e.g. commercial, scientific), and included re-exports because this information was rarely known for the seizures data. We analysed all reported legal imports to the US and the EU, arriving directly from the country of origin or imported indirectly via one or more interim country, aggregated across all exporting countries. For the EU, to ensure consistency between EU-TWIX and the CITES Trade Database, data records were filtered to exclude those involving trade between Member States (i.e. all reported trade analysed was that entering the EU from outside its borders).

For US seizures, data originating from the LEMIS database are reported as part of the US annual reports to CITES. We extracted these data from the CITES Trade Database by filtering the shipment records for those where the importing Party was the US and the source of the shipment was coded as "confiscated or seized" ("I") as per CITES Resolution 12.3 (Rev. CoP17). Data were extracted for the same time-period (1975-2014) as the reported trade volumes, though most time-series began later, with LEMIS becoming operational in 1983 (no LEMIS time-series began before 1982). For EU seizures, the EU-TWIX database is the only EU-wide wildlife seizures database. At the time of analysis, the database held over 50,000 records from the 28 EU Member States, covering the period 2000-2016, reported by a range of CITES enforcement authorities, including Customs, police, wildlife inspection services, and so forth. The EU-TWIX dataset was downloaded in July 2016, following permission granted by the EU Member States. We analysed the period 2005 to 2014, representing the longest period over which relatively complete seizure information was available in the database at the time of analysis. We used data from all 28 EU Member States from the point that they acceded. The US and the EU are themselves major end markets for illegal commodities. Specifically, for US seizures, 28% of the time-series originated in a known country other than the US and were being re-exported from the US. For EU-TWIX seizures, 5% originated outside the EU and were not destined for the EU.

All outputs from the EU-TWIX database were checked for consistency in taxonomic names and standardised according to CITES standard nomenclature. To reconcile taxonomy, any synonyms reported in EU-TWIX were changed to current CITES-accepted names based on the list extracted from Species+ (speciesplus.net) and the CITES Checklist. If the taxon name could not be reconciled to an accepted CITES species, the seizure record was removed from the dataset (1.4% of EU-TWIX records). LEMIS data were already standardised following CITES nomenclature. We focussed on taxa that were identified to species level, to ensure that we were not confounding differing species in reported trade and seizures, as well as to ensure that our results could be related back to the sustainability of individual species.

Records were aggregated to unique taxon, product and unit combinations ('taxon-products') to yield 18,990 time-series of reported trade volumes into the US and 7,730 time-series of seizure volumes (1982 to 2014); and 17,137 time-series of reported trade volumes into the EU and 3,577 time-series of seizure volumes (2005-2014). As the seizures data were more intermittent, and our focus was on testing for consistency of relationships, both positive and negative, between reported and seized trade, we focussed on taxon-products for which there were relatively consistent seizures in multiple years (i.e. which were frequently interdicted) and for which seizure volumes were reasonably high (i.e. which were information rich). We therefore ranked the time-series of seizures for the US and the EU based on the number of years in which seizures were made (i.e. non-zero records), and retained the minimum number of time-series that accounted for 50% or more of the total number of seizures (across all taxon-products retained). For LEMIS this equated to all time-series with 16 or more non-zero data years between 1982 (or start of time-series) and 2014; for EU-TWIX, this corresponded to time-series for which there were 7 or more non-zero data years between 2005 and 2014. In total, there were 163 EU-TWIX time-series and 301 LEMIS time-series which met these criteria.

We removed products from the analysis for which we could not be sure of the number or amount of wildlife they contained and which were hence unable to be compared between time-series (medicines, powder, waxes, and derivatives). As most plant species were listed as medicines or other derivatives, and/or only identified to family level in one or several of the databases, this meant that no plant species remained in the analysis.

We then reviewed the time-series of trade volumes for wildlife species/subspecies and product types that are traded equivalently (i.e. are essentially the same good in the destination market) and could therefore be combined in a single collective time-series; for example, leather goods from *Caiman crocodilus* and its subspecies *Caiman crocodilus crocodilus* (for full details see Appendix S1 and Table S1). To ensure the robustness of the data for analysis, we took steps to exclude taxa for which clear external impacts or differences in the legislative environment may have introduced biases. Species that had been transferred between the CITES Appendices during the analysis period were excluded, to prevent confounding effects on the association between reported trade volumes and seizures. Similarly, taxa that were split-listed (with populations on both Appendix I and II) were similarly excluded; thus, some iconic species such as African elephant (*Loxodonta africana*) were not included in the analysis. Finally, sub-species that were listed in a different Appendix to their parent species were also excluded. One exception to the above procedures was caviar (from *Acipenseriformes*), given the importance of the product and its interchangeability.

Despite retaining time-series with relatively frequent seizures, there were still some with low mean seizure volumes or few years of legally-reported trade. To ensure that the analysis was conservative and that we did not over-interpret based on low-volume species, we lastly removed time-series with a mean number of seized products per year of less than five, or fewer than three years of legal transactions. After all quality control processes, we were left with 28 time-series for the US and 20 for the EU, representing combined taxon-products which were equivalent among databases, identified to species or genus level, and having relatively high volumes and frequencies of both trade and seizures.

*Modelling approach for relating reported trade to law enforcement seizures*

In principle, there may be many idiosyncratic relationships between legal and illegal wildlife trade for different products and species, which can be dynamic through both space and time depending upon the political, cultural, enforcement, and economic contexts. In practice, we could anticipate a positive, negative, or no relationship between reported legal trade volumes and seizure volumes for any destination market, with the null expectation being 'no relationship'.

Specifically, our hierarchical Bayesian regression modelling framework had the seizure volume as the response variable, with the legal trade volume, enforcement effort, and (for the EU) reporting level as co-variates. Our analysis considered the US and the EU as separate destination markets (i.e. a separate model was fitted for each). For each destination market, the model assumed that the number of reported seizures $s_{j,t}$ for taxon-product $j$ in year $t$ had a negative binomial distribution such that:

$$s_{j,t} \sim NB(\mu_{j,t}, \phi_j)$$

where the parameter $\mu_{j,t}$ is the mean for taxon-product $j$ in year $t$, and $\phi_j$ is an overdispersion parameter such that the variance is $\frac{\mu_{j,y}^2}{\phi_j}$. The mean $\mu_{j,t}$ is a function of the reported legal trade volume, $l_t$, the enforcement 'effort', $e_t$, as measured by a proxy variable, and the reporting rate, $r_t$:

$$\mu_{j,t} = exp(\alpha_j + \beta_j l_t + \gamma_j e_t) r_t$$

with $\alpha_j$ an intercept, $\beta_j$ the slope parameter relating the legal trade volume to the seizure volume (the key parameter of interest), and $\gamma_j$ a parameter that controls for enforcement effort with respect to seizures. The hierarchical component of the model is derived from the fact that the slopes $\beta_j$ for individual taxon-products in the same destination market are were drawn from a normal distribution such that:

$$\beta_j \sim Normal(\beta, \sigma^2)$$

where $\beta$ is the mean slope across taxon-products and $\sigma^2$ is the variance.

The posterior estimate of $\beta$ therefore represents the mean of the overall relationship between legal trade volume and seizures across all taxon-products in a destination market. We assessed differences among taxa, CITES Appendix, and product type by testing for differences in posterior distributions.

We included the total number of seizures (as opposed to volumes) across all taxon-products in each year in the US or the EU to account for variation in enforcement effort, $e_t$, over time. While the seizures of specific individual taxon-products may fluctuate substantially due to changes in demand or supply, the total number of seizures is more likely to reflect enforcement effort than total illegal trade volume, notwithstanding any substantial changes in (illegal) shipments over time summed across *all* illegal wildlife trade goods. Thus, we assume that this proxy reflects on the ground (border) enforcement effort reasonably well. We check this assumption in the United States through comparing results to an alternative proxy in which the effort variable is the total number of containers inspected each year. We selected the total number of seizures as the primary effort proxy because it was almost twice as long in terms of number of years, and hence enabled us to use the full length of the US time-series. See Appendix S2 for details.

In general, our model is somewhat similar to an analysis of elephant ivory trade(Underwood et al. 2013), except that the effort enters additively as a covariate rather than a logit-transformed probabilistic multiplication factor. In a similar manner to slope parameters linking legal trade and seizures, effort parameters $\gamma_j$ relating the increase in seizures for each taxon-product $j$ as enforcement effort increases were drawn from a normal distribution, enabling a separate relationship to be characterized for each taxon-product.

To account for differences in the EU Member States of reporting rates of seizures to CITES, $r_t$, we included a covariate consisting of the proportion of countries in each year that had provided a complete record of seizures to EU-TWIX at the time of the analysis (>95% for all years except 2014), weighted by the mean relative proportion of EU-TWIX seizures ascribed to that country (Table S2). We assumed that countries that had partially reported seizures (i.e. did not provide information on all seizures during a whole year) did not contribute, and only included those countries that reported completely. For the US, the US Fish and Wildlife Service Office of Law Enforcement documents all known CITES wildlife trade, legal and illegal, within the LEMIS database (except plant species which are less commonly documented). Given that the reporting rate of CITES-related seizures by the USFWS in annual reports to CITES is very high, and the time elapsed since the last year of the analysis (2014), in the absence of additional information we assumed that the proportional reporting rate each year in the US was one.

*Model fitting*

All models were fitted within a Bayesian modelling framework using a sample-based Hamiltonian Markov Chain Monte Carlo algorithm (Stan Development Team 2015). We tested the estimation algorithm on simulated data prior to being fitted to time-series to ensure that they could recapture known parameters. The algorithm randomly iterates through potential parameter combinations to find combinations that yield high posterior probability. Four implementations of the algorithm (i.e. ‘chains’) were run with 4000 iterations each. Each chain contained 2000 warmup iterations which helps initialize the chains. No thinning of chains was done as the Hamiltonian algorithm produced chains with little correlation between iterates. We used broad uninformative priors for the model parameters (normal priors with means of zero and variances of 1000 for mean parameters, and Cauchy priors with means of zero and variances of five for variance parameters).

Legal trade values and effort proxies (total number of seizures & number of shipments inspected; see Appendix S2) were z-transformed prior to being included in the model in order to aid convergence of the algorithm. Some models experienced convergence issues when there were very large 'spikes' in individual year trade volumes. To handle these large statistical outliers in the legal and illegal time-series, we applied an influence function from robust statistics(Wilcox 2005) that trimmed outlier values such that no data point was larger than three times the median absolute deviation from the median time-series value, while preserving the relative magnitudes of the original outliers. Post-fitting inspection for potential model convergence issues included examining the scale reduction factor, plotting the model fit and the behaviour of each chain, and bivariate plots of variables and the linear predictor. Furthermore, we calculated an $R^2$ for each model as a simple measure of variance explained, according to the means of the posterior distributions for the model parameters.

All models were fitted in the statistical software R (R Core Team 2016) using the *rStan* package (Rstan Team 2018).

**Appendix S2: Combination of products and taxa**

We combined several taxa together where they would likely be traded interchangeably as the same product. For example, *Caiman crocodilus* was combined with its subspecies *Caiman crocodilus crocodilus* and *Caiman crocodilus fuscus* when the product type was leather goods. Leather products were reported as 'large' or 'small' leather products, leather items, garments, or shoes; we combined these to create a single leather products time-series for relevant taxa. Ivory was recorded using different terms such as carvings, tusks or ivory products, and these varied between databases, so were combined treated equivalently. Trophies and skulls were also combined, as nomenclature may vary between databases and enforcement organisations but represent similar products. Three species (*Acipenser baerii*, *Acipenser sturio*, and *Huso huso*) that produced caviar were combined as this product is likely traded interchangeably; this special case is the only instance in which Appendix II (*Acipenser baerii*, *Huso huso*) and Appendix I (*Acipenser sturio*) species were combined, though note that none of these species changed annex during the period of analysis. For full details of all taxa and products that were combined, see Table S1. As indicated in the main text, we also removed commodities such as medicines, powders or derivatives since it was unclear what volume of the associated taxa was contained in the seizure; often only trace amounts.

**Appendix S2: Evaluation of enforcement effort proxies**

The robustness analysis on the proxies for enforcement effort was conducted by using the total number of containers inspected by the US Fish & Wildlife Service (UNODC 2016) as an alternative proxy to total seizures across all taxa (Figure S3). As container inspection data were only available back to 1998, this also necessitated re-fitting models to shortened time series (beginning in 1998). For comparison, we also refit using the original effort proxy (total number of seizures across all taxa) but shortened to begin only in 1998 (rather than 1982) (Figure S4).

The overall meta-analytic median was not significantly different between using container inspections and the original effort proxy, suggesting that broad trends remained similar. However, the meta-analytic medians for both the container inspections and the shortened original effort proxy were non-significant (though still positive). Importantly, given that the time-series were much shorter than the full data, some taxon-products (container inspections and an equivalently short time-series of total seizures) showed differences in result when compared to the model fitted to the whole time-series. This suggests that short time-series can show results that may be different from those over longer time-scales, perhaps due to longer time-series containing distinct patterns over shorter periods. In summary, we found that: (i) both effort proxies gave a positive meta-analytic median for the relationship between legal trade and seizure volumes into the US, though this was only significant for the full-length original effort proxy (total number of seizures across all CITES-listed taxon-products); (ii) while meta-analytic medians were not significantly different from one another, there were some differences in individual taxon-product slopes between effort proxies, suggesting some differences in pattern between long and short time-series; this is reinforced by repeating the analysis using a shortened original effort proxy that begins in 1998, the same year as the container inspections.

**Table S1: Taxa and products combined (due to being traded near-equivalently in destination markets)**

| Taxon 1 | Taxon 2 | Taxon 3 | Unit (where relevant) | Terms† | | | | |
|---|---|---|---|---|---|---|---|---|
| Loxodonta africana | | | | CAR | IVP | TUS | | |
| Loxodonta africana | | | (kg) | CAR | TUS | | | |
| Python reticulatus | | | | LPS | LPL | | | |
| Python sebae | | | | LPS | LPL | | | |
| Naja naja | | | | LPS | LPL | | | |
| Chelonia mydas | | | | BOD | CAP | | | |
| Python molurus | Python molurus molurus | | | LPS | LPL | | | |
| Python molurus | Python molurus molurus | | | SKI | | | | |
| Eretmochelys imbricata | | | | BOD | CAP | | | |
| Chelonia mydas | | | | BOD | CAP | | | |
| Loxodonta africana | | | | IVC | TUS | CAR | TEE | IVP |
| Caiman crocodilus | Caiman crocodilus crocodilus | Caiman crocodilus fuscus | | LPS | SHO | LPL | SKO | |
| Crocodylus niloticus | | | | LPS | LPL | SKO | | |
| Python sebae | | | | LPS | SHO | | | |
| Varanus niloticus | | | | LPS | SHO | | | |
| Alligator mississippiensis | | | | LPS | LPL | SHO | | |
| Python reticulatus | | | | LPS | SHO | GAR | LPL | SKO |
| Elephas maximus | | | | IVC | TUS | TEE | IVP | CAR |
| Caiman crocodilus | Caiman crocodilus crocodilus | | | BOD | | | | |
| Varanus salvator | | | | LPS | SHO | | | |
| Hippopus hippopus | | | | SHE | CAR | | | |
| Caiman crocodilus | Caiman crocodilus fuscus | | | SKI | | | | |
| Odobenus rosmarus | | | | TUS | CAR | | | |
| Loxodonta africana | | | | LPS | SHO | SKO | | |
| Ptyas mucosus | | | | LPS | SHO | | | |
| Hippopotamus amphibius | | | | TEE | TUS | | | |
| Leopardus pardalis | | | | GAR | LPS | | | |
| Python bivittatus | | | | LPS | SHO | | | |
| Crocodylus moreletii | | | | LPS | SHO | | | |
| Tupinambis teguixin | | | | LPS | SHO | | | |
| Acipenser baerii | Acipenser sturio | Huso huso | | CAV | | | | |

† BOD = bodies; CAP = carapaces; CAR = carvings; GAR = garments; IVC = ivory carvings; IVP = ivory pieces; LPL = large leather products; LPS = small leather products; SHE = shells; SHO = pairs of shoes; SKI = skins; SKO = leather items; TEE = teeth; TUS = tusks.

**Table S2: Proportion of EU countries in each year that had given a complete report on seizures to EU-TWIX (weighted by total seizures). Source: EU-TWIX database, 2016 (https://www.eu-twix.org/).**

| **Year** | **Proportion reported (weighted by proportion of total seizures)** |
| --- | --- |
| 2005 | 0.998 |
| 2006 | 0.998 |
| 2007 | 0.989 |
| 2008 | 0.991 |
| 2009 | 0.953 |
| 2010 | 0.968 |
| 2011 | 0.969 |
| 2012 | 0.985 |
| 2013 | 0.950 |
| 2014 | 0.834 |

**Table S3: Total number of seizures per year (across all taxa and products)**

| Year | Number of seizures in LEMIS (US) | Number of seizures in EU-TWIX (EU) |
|---|---|---|
| 1982 | 605 | |
| 1983 | 781 | |
| 1984 | 659 | |
| 1985 | 920 | |
| 1986 | 1032 | |
| 1987 | 1091 | |
| 1988 | 1168 | |
| 1989 | 1450 | |
| 1990 | 1892 | |
| 1991 | 1777 | |
| 1992 | 1569 | |
| 1993 | 1523 | |
| 1994 | 1356 | |
| 1995 | 1274 | |
| 1996 | 1207 | |
| 1997 | 1177 | |
| 1998 | 1467 | |
| 1999 | 1528 | |
| 2000 | 1371 | |
| 2001 | 1251 | |
| 2002 | 1256 | |
| 2003 | 1318 | |
| 2004 | 1404 | |
| 2005 | 1561 | 1282 |
| 2006 | 1604 | 1658 |
| 2007 | 1527 | 1270 |
| 2008 | 2613 | 1306 |
| 2009 | 2758 | 1445 |
| 2010 | 2559 | 1305 |
| 2011 | 2578 | 1360 |
| 2012 | 2422 | 1476 |
| 2013 | 2154 | 1644 |

| | | |
|---|---|---|
| 2014 | 1977 | 1311 |

**Table S4: Number of container shipments inspected across the US by the US Fish and Wildlife Service. Source: World Wise database** (UNODC 2016)**.**

| **Year** | **Number of shipments inspected** |
|---|---:|
| 1998 | 86,409 |
| 1999 | 75,252 |
| 2000 | 111,296 |
| 2001 | 116,535 |
| 2002 | 123,720 |
| 2003 | 138,754 |
| 2004 | 157,617 |
| 2005 | 171,874 |
| 2006 | 183,247 |
| 2007 | 187,670 |
| 2008 | 186,959 |
| 2009 | 176,798 |
| 2010 | 178,341 |
| 2011 | 183,063 |
| 2012 | 184,564 |
| 2013 | 184,824 |
| 2014 | 180,463 |

**Supplementary Figures**

**Figure S1 (see separate .pdf file):** Time-series plots, model fits, and slope estimates for US time-series. Left-hand column shows legal trade volume (black circles and lines) and seizure volume (red squares and lines). Data have been processed using a robust statistics approach as described in the Methods. Central column shows the posterior median (black line) and mean (red line) estimates of seizure volume, along with the 95% credible intervals (dashed line). Black circles represent seizures. Right-hand column shows median model estimate and 95% credible intervals for slope ($\beta$) parameter representing the relationship between legal trade volume and seizures volume, along with log-scale partial residuals. Enforcement effort and reporting proxies also included as co-variates. CITES data are importer-reported and include all sources except source “I”.

**Figure S2 (see separate .pdf file):** Time-series plots, model fits, and slope estimates for EU time-series. Left-hand column shows legal trade volume (black squares and lines) and seizure volume (red squares and lines). Data have been processed using a robust statistics approach as described in the Methods. Central column shows the posterior median (black line) and mean (red line) estimates of seizure volume, along with the 95% credible intervals (dashed line). Black circles represent seizures. Right-hand column shows median model estimate and 95% credible intervals for slope ($\beta$) parameter representing the relationship between legal trade volume and seizures volume, along with log-scale partial residuals. Enforcement effort and reporting proxies also included as co-variates. CITES data are importer-reported and include all sources except source “I”.

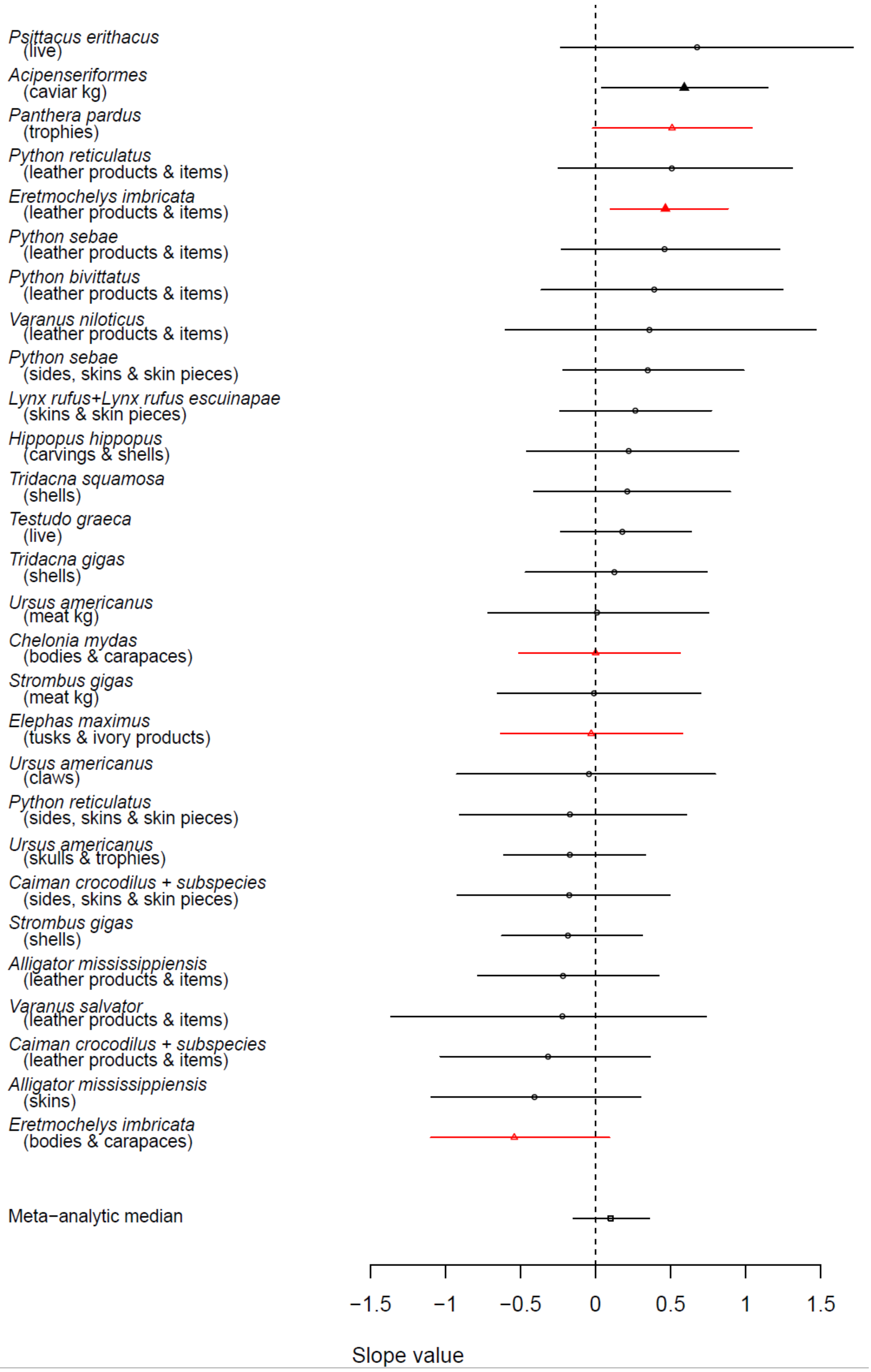

Psittacus erithacus (live)
Acipenseriformes (caviar kg)
Panthera pardus (trophies)
Python reticulatus (leather products & items)
Eretmochelys imbricata (leather products & items)
Python sebae (leather products & items)
Python bivittatus (leather products & items)
Varanus niloticus (leather products & items)
Python sebae (sides, skins & skin pieces)
Lynx rufus+Lynx rufus escuinapae (skins & skin pieces)
Hippopus hippopus (carvings & shells)
Tridacna squamosa (shells)
Testudo graeca (live)
Tridacna gigas (shells)
Ursus americanus (meat kg)
Chelonia mydas (bodies & carapaces)
Strombus gigas (meat kg)
Elephas maximus (tusks & ivory products)
Ursus americanus (claws)
Python reticulatus (sides, skins & skin pieces)
Ursus americanus (skulls & trophies)
Caiman crocodilus + subspecies (sides, skins & skin pieces)
Strombus gigas (shells)
Alligator mississippiensis (leather products & items)
Varanus salvator (leather products & items)
Caiman crocodilus + subspecies (leather products & items)
Alligator mississippiensis (skins)
Eretmochelys imbricata (bodies & carapaces)
Meta−analytic median
−1.5
−1
−0.5
0
0.5
1
1.5
Slope value

**Figure S3:** *Strength of the relationship between the legal trade volume and seizure volume over time for the US, using the total number of containers as an effort proxy. Note that this resulted in shorter time-series (beginning in 1998 instead of early 1980s for most taxa). Values are the slope parameter from a hierarchical Bayesian regression model relating reported trade as a predictor of seizure volume (with units increase in seizures per unit change in legal trade). A positive value indicates a positive relationship between reported trade and seizure volumes, while a negative value indicates an inverse relationship. The meta-analytic median represents the hierarchical slope parameter value. Symbols indicate posterior Bayesian median model estimates, while lines indicate 95% credible intervals. Filled symbols indicate a value that is significantly different from zero while open circles indicate a non-significant relationship. Red triangles indicate CITES Appendix I species; all other species CITES Appendix II during the analysis period. Note that Acipenseriformes (black triangle) includes one Appendix I and two Appendix II species. Taxa ordered by effect size.*

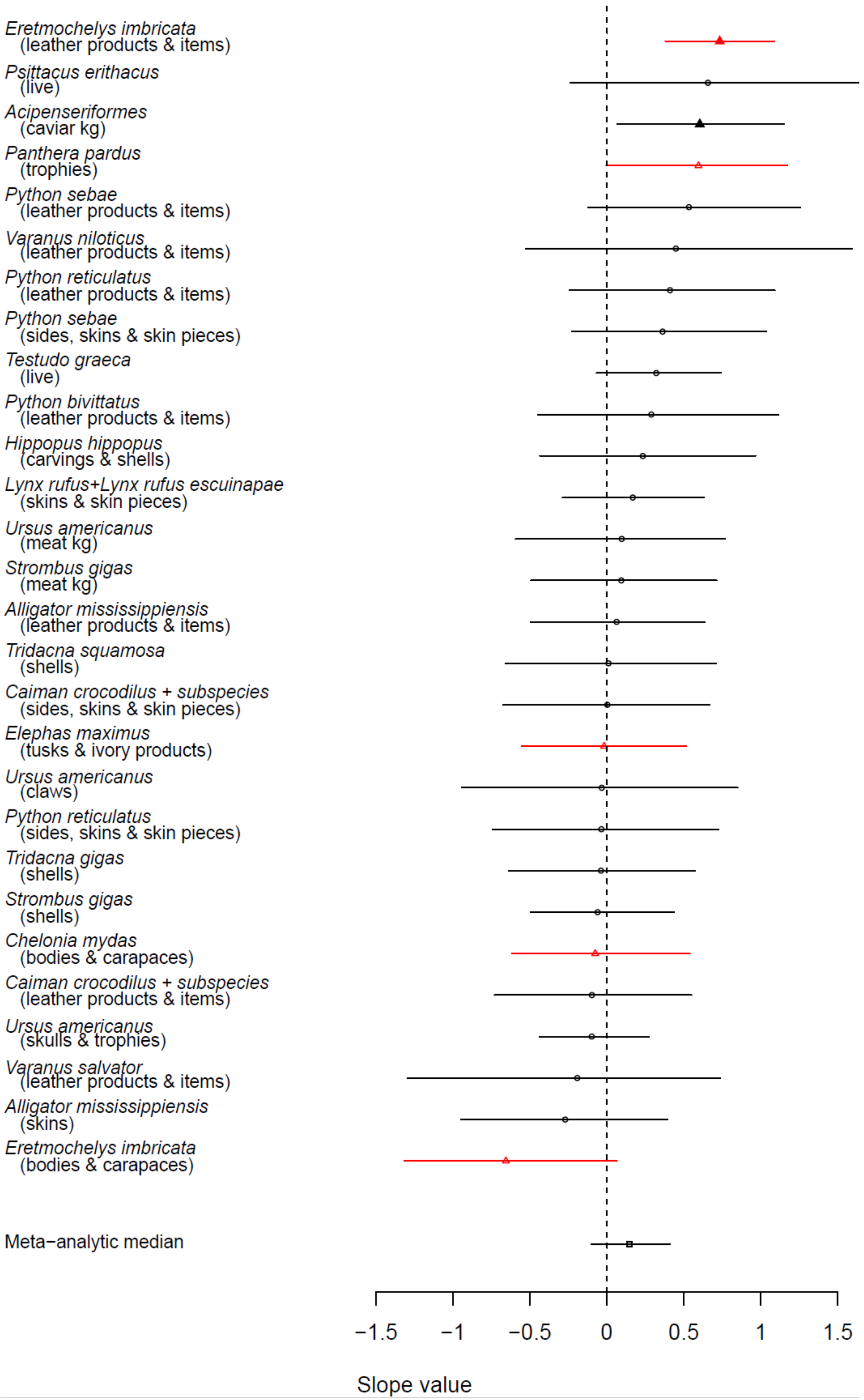
Eretmochelys imbricata (leather products & items)
Psittacus erithacus (live)
Acipenseriformes (caviar kg)
Panthera pardus (trophies)
Python sebae (leather products & items)
Varanus niloticus (leather products & items)
Python reticulatus (leather products & items)
Python sebae (sides, skins & skin pieces)
Testudo graeca (live)
Python bivittatus (leather products & items)
Hippopus hippopus (carvings & shells)
Lynx rufus+Lynx rufus escuinapae (skins & skin pieces)
Ursus americanus (meat kg)
Strombus gigas (meat kg)
Alligator mississippiensis (leather products & items)
Tridacna squamosa (shells)
Caiman crocodilus + subspecies (sides, skins & skin pieces)
Elephas maximus (tusks & ivory products)
Ursus americanus (claws)
Python reticulatus (sides, skins & skin pieces)
Tridacna gigas (shells)
Strombus gigas (shells)
Chelonia mydas (bodies & carapaces)
Caiman crocodilus + subspecies (leather products & items)
Ursus americanus (skulls & trophies)
Varanus salvator (leather products & items)
Alligator mississippiensis (skins)
Eretmochelys imbricata (bodies & carapaces)
Meta−analytic median
−1.5
−1
−0.5
0
0.5
1
1.5
Slope value

**Figure S4:** *Strength of the relationship between the legal trade volume and seizure volume over time for the US, using a (shortened) total number of seizures as an effort proxy. Note that this resulted in shorter time-series (beginning in 1998 instead of early 1980s for most taxa). Values are the slope parameter from a hierarchical Bayesian regression model relating reported trade as a predictor of seizure volume (with units increase in seizures per unit change in legal trade). A positive value indicates a positive relationship between reported trade and seizure volumes, while a negative value indicates an inverse relationship. The meta-analytic median represents the hierarchical slope parameter value. Symbols indicate posterior Bayesian median model estimates, while lines indicate 95% credible intervals. Filled symbols indicate a value that is significantly different from zero while open circles indicate a non-significant relationship. Red triangles indicate CITES Appendix I species; all other species CITES Appendix II during the analysis period. Note that Acipenseriformes (black triangle) includes one Appendix I and two Appendix II species. Taxa ordered by effect size.*
