## Supplementary figures and images for "Evaluating the relationships between the legal and illegal international wildlife trades"

### Figure S1

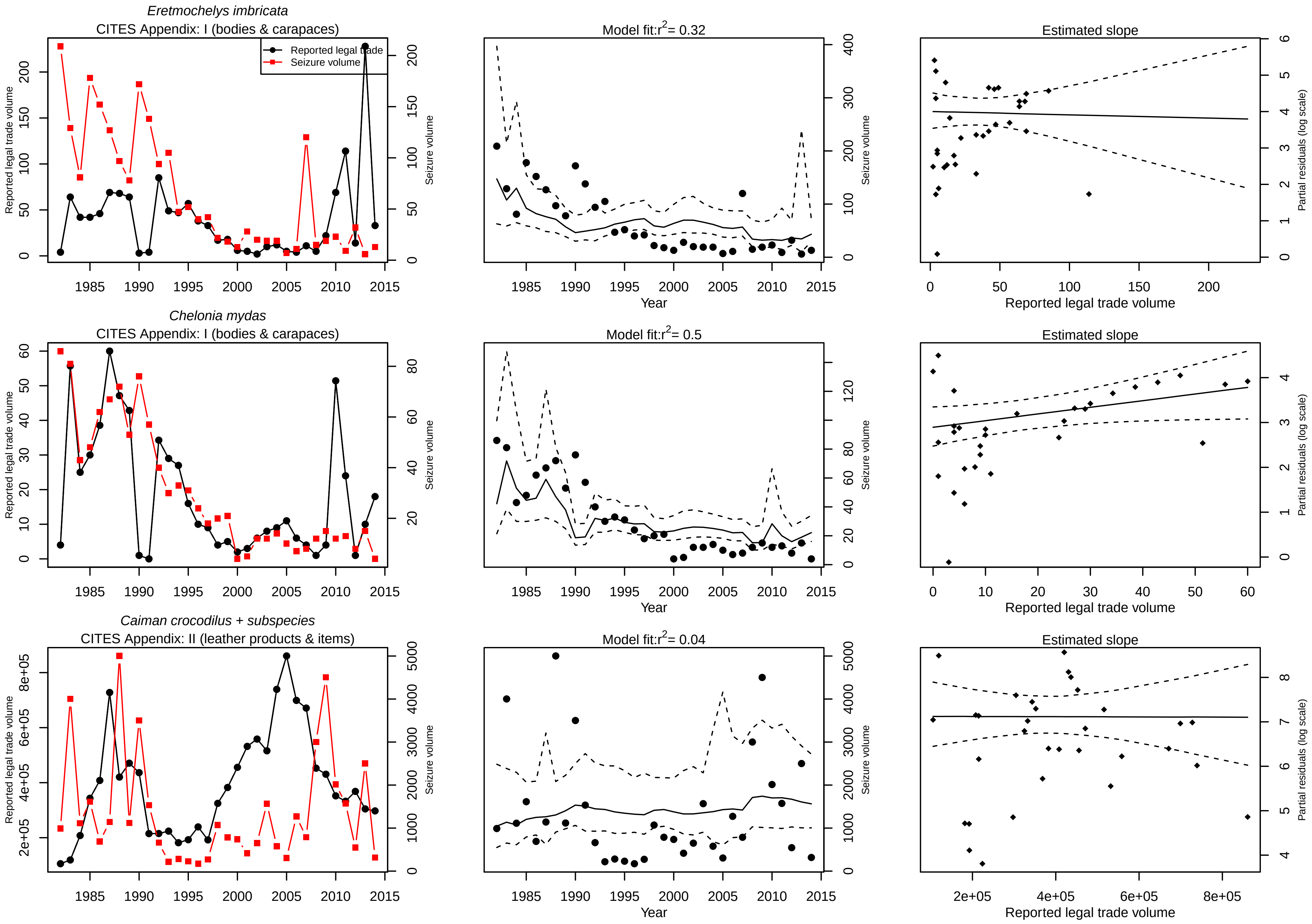

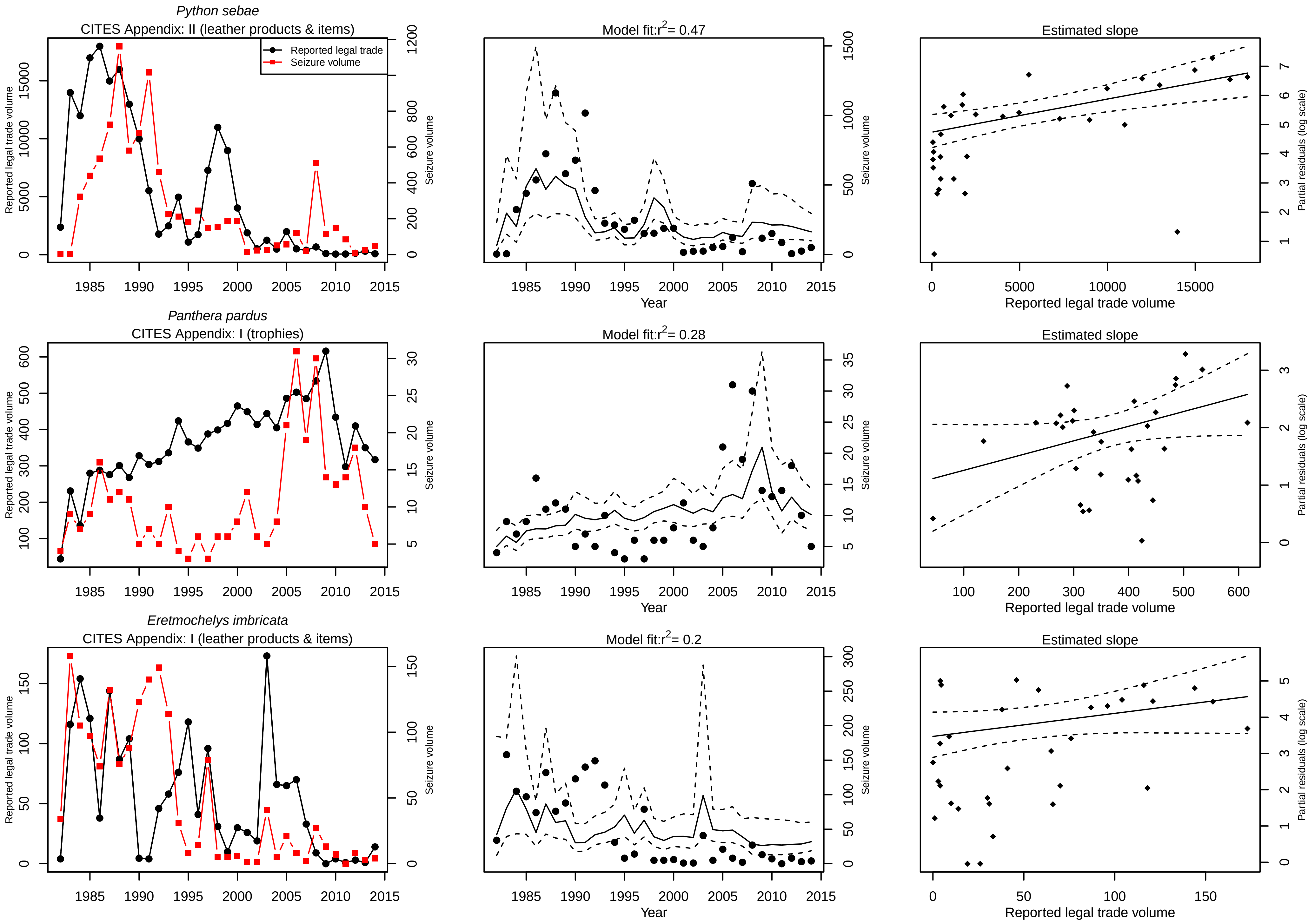

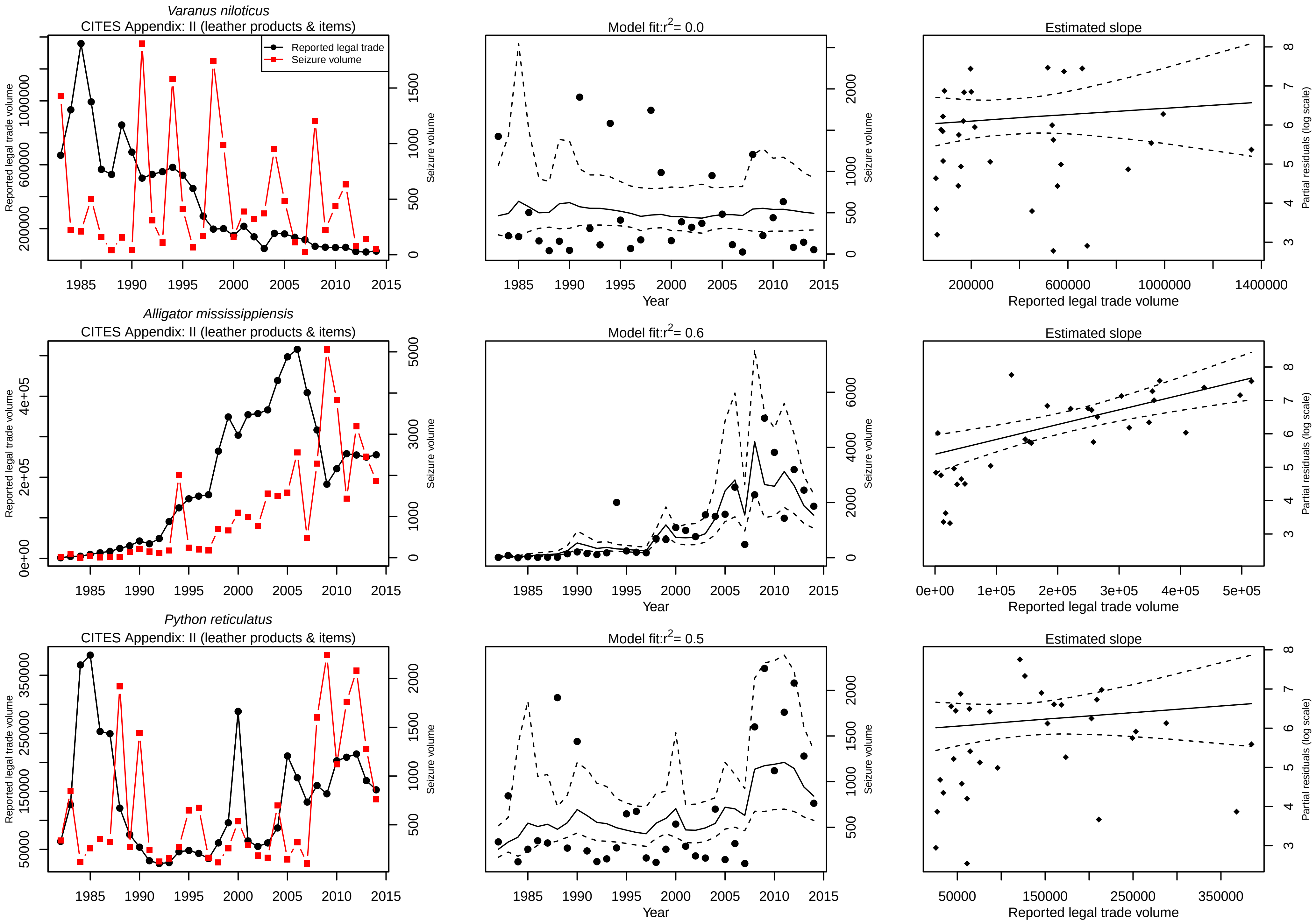

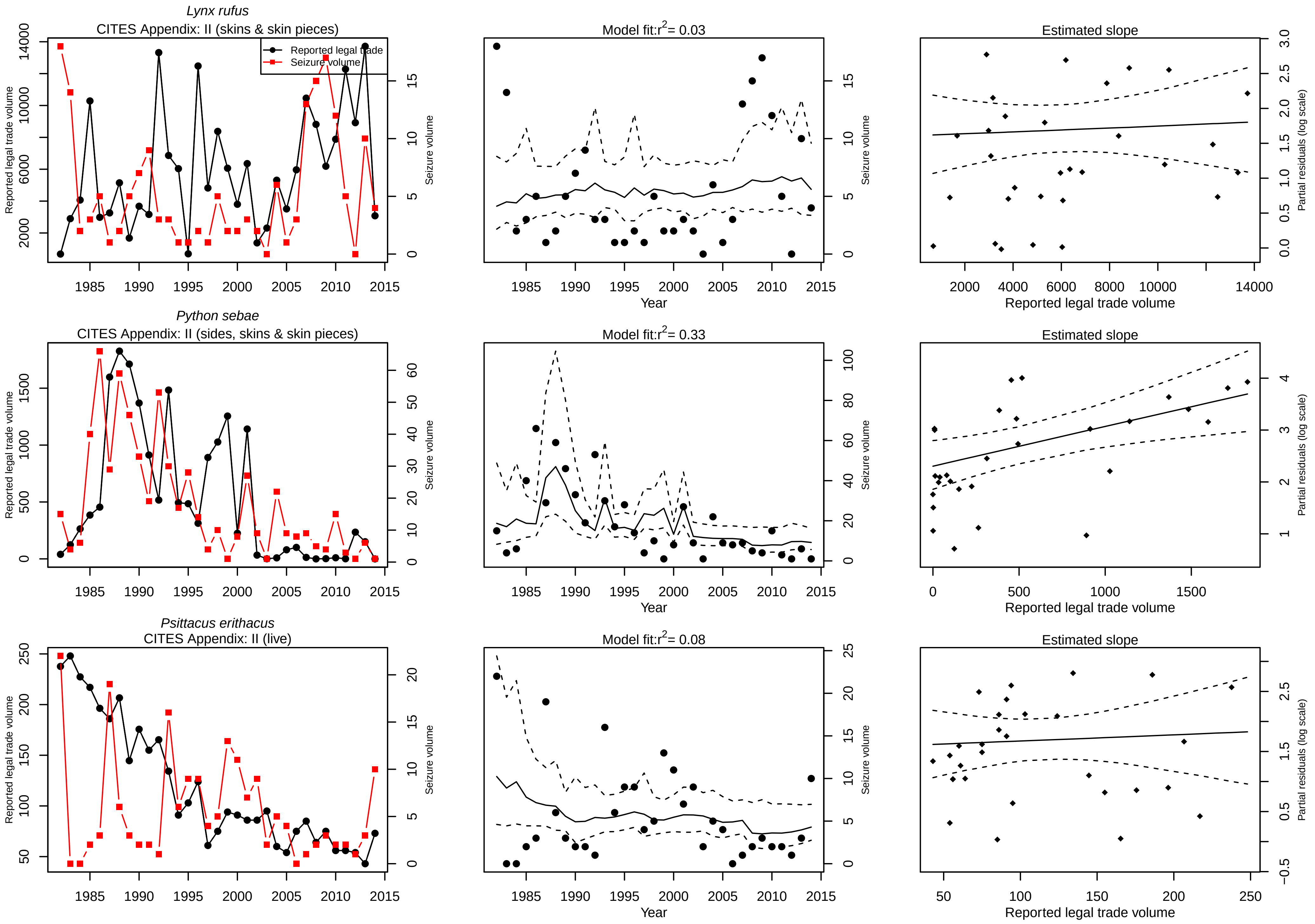

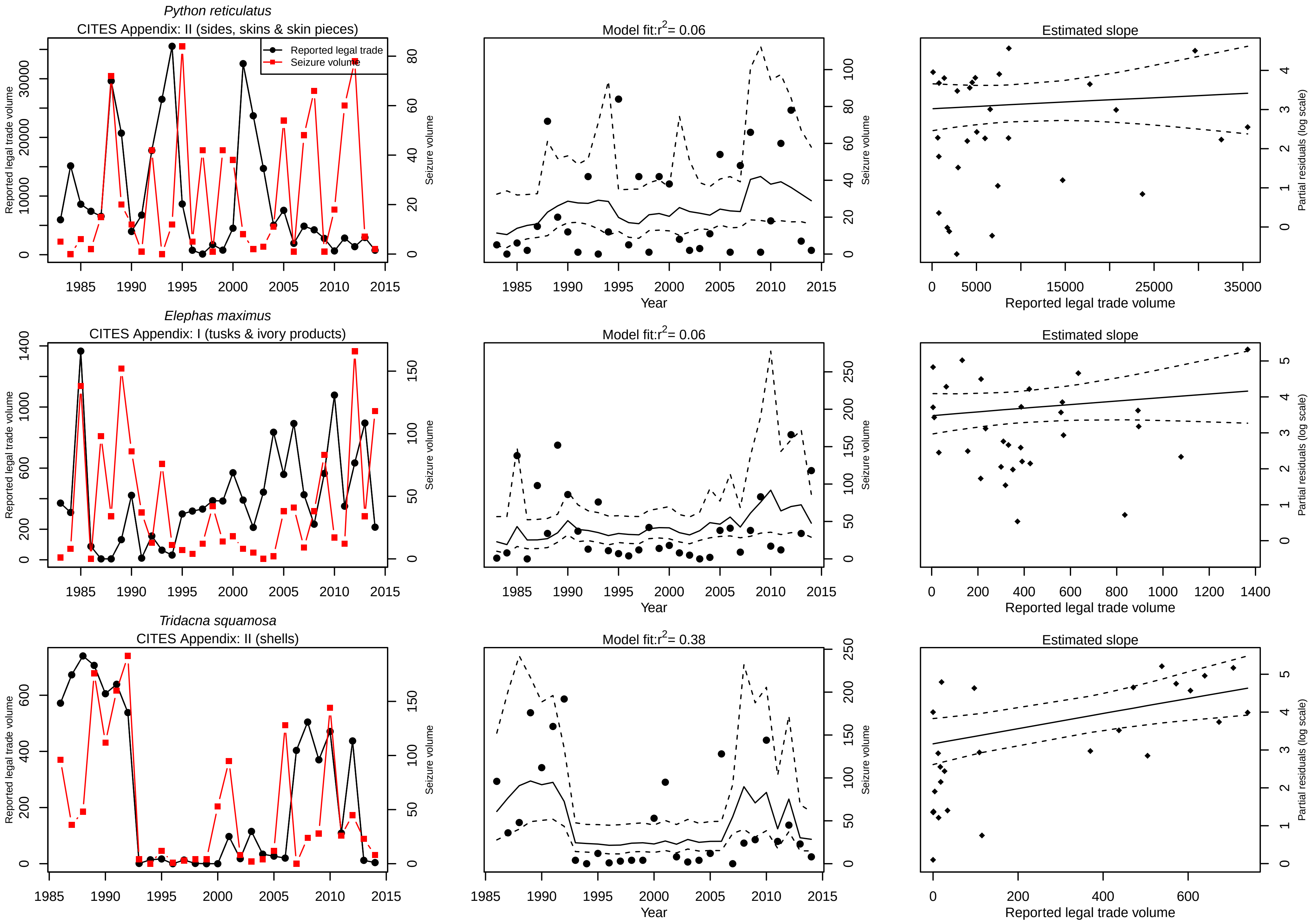

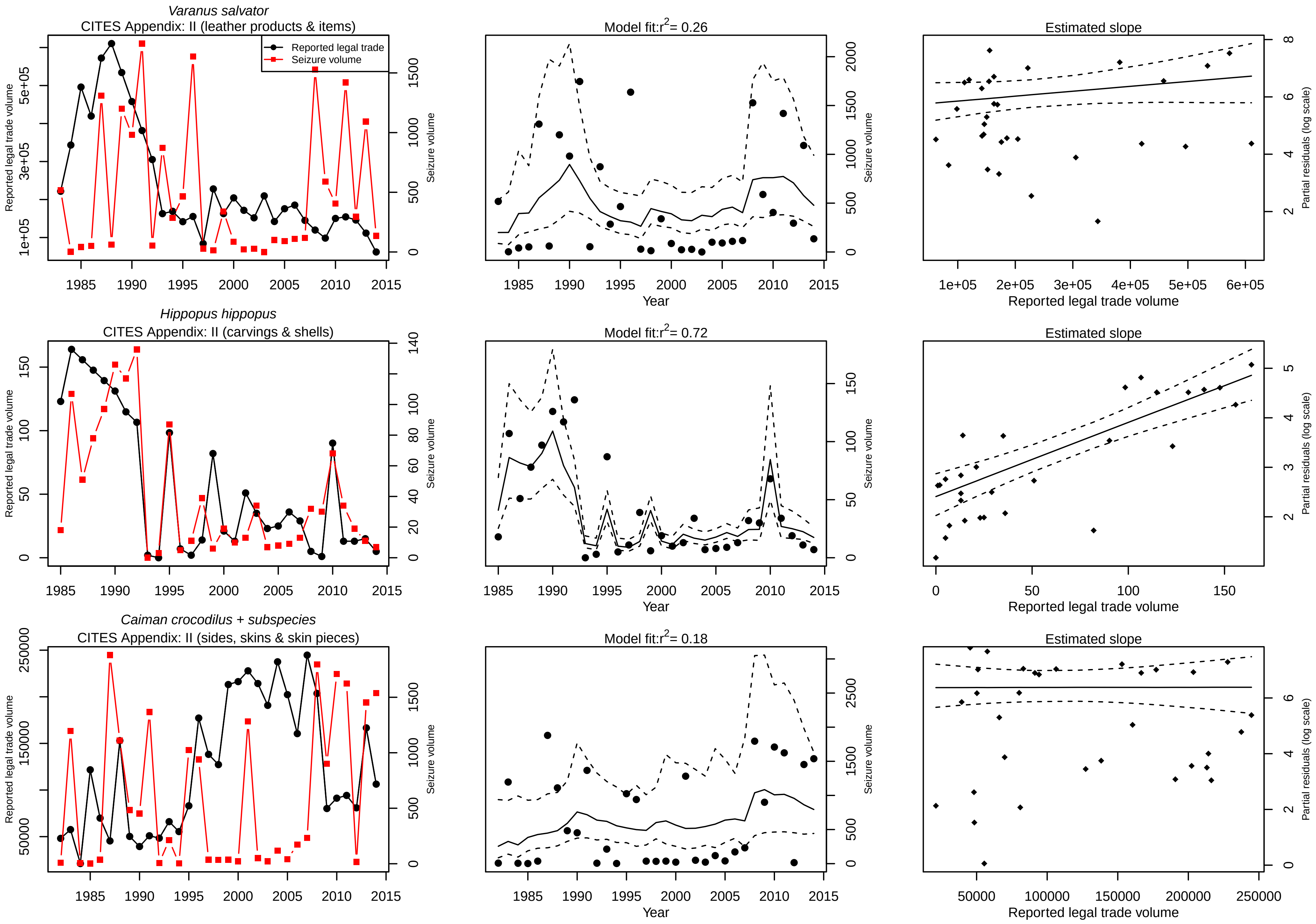

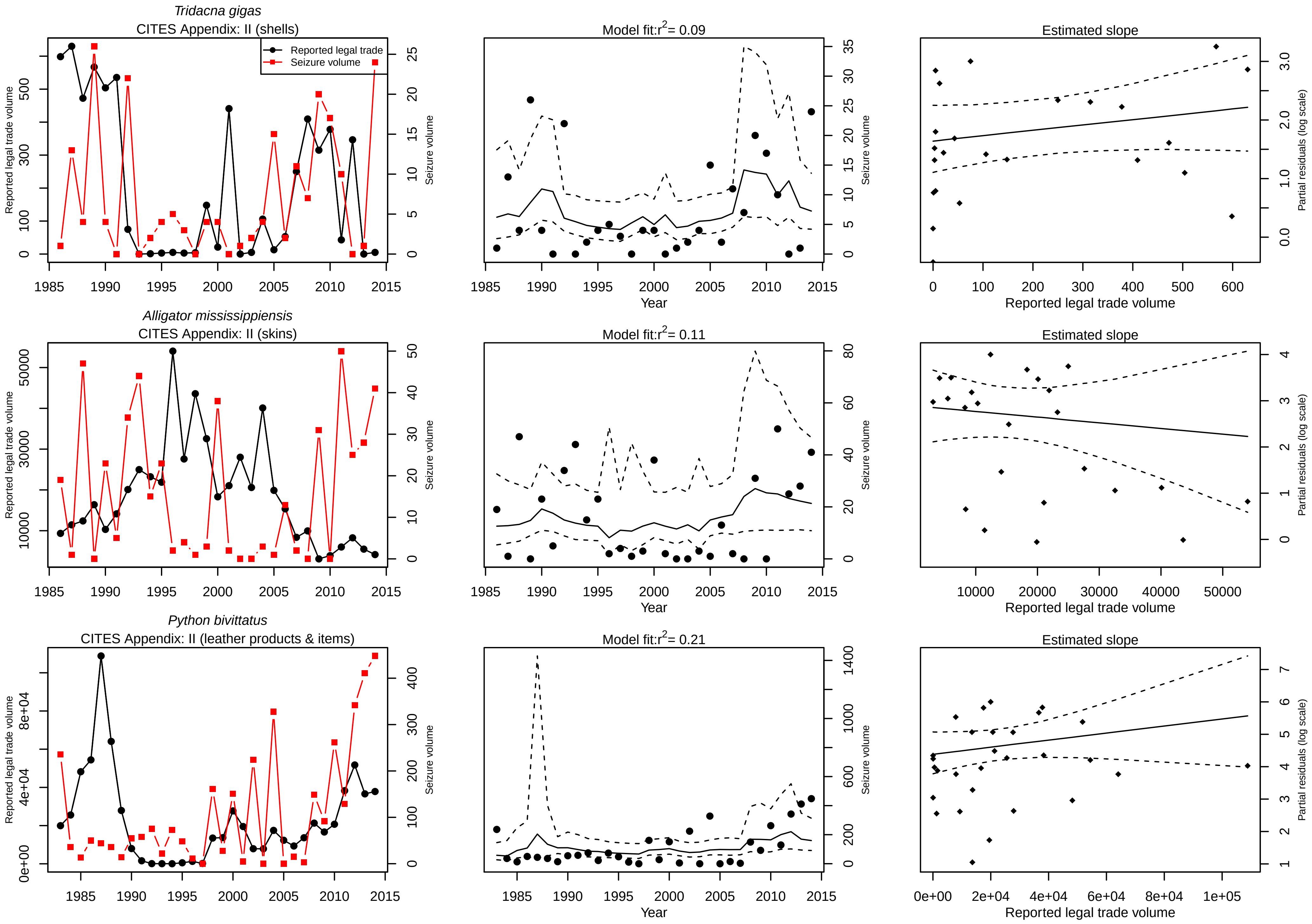

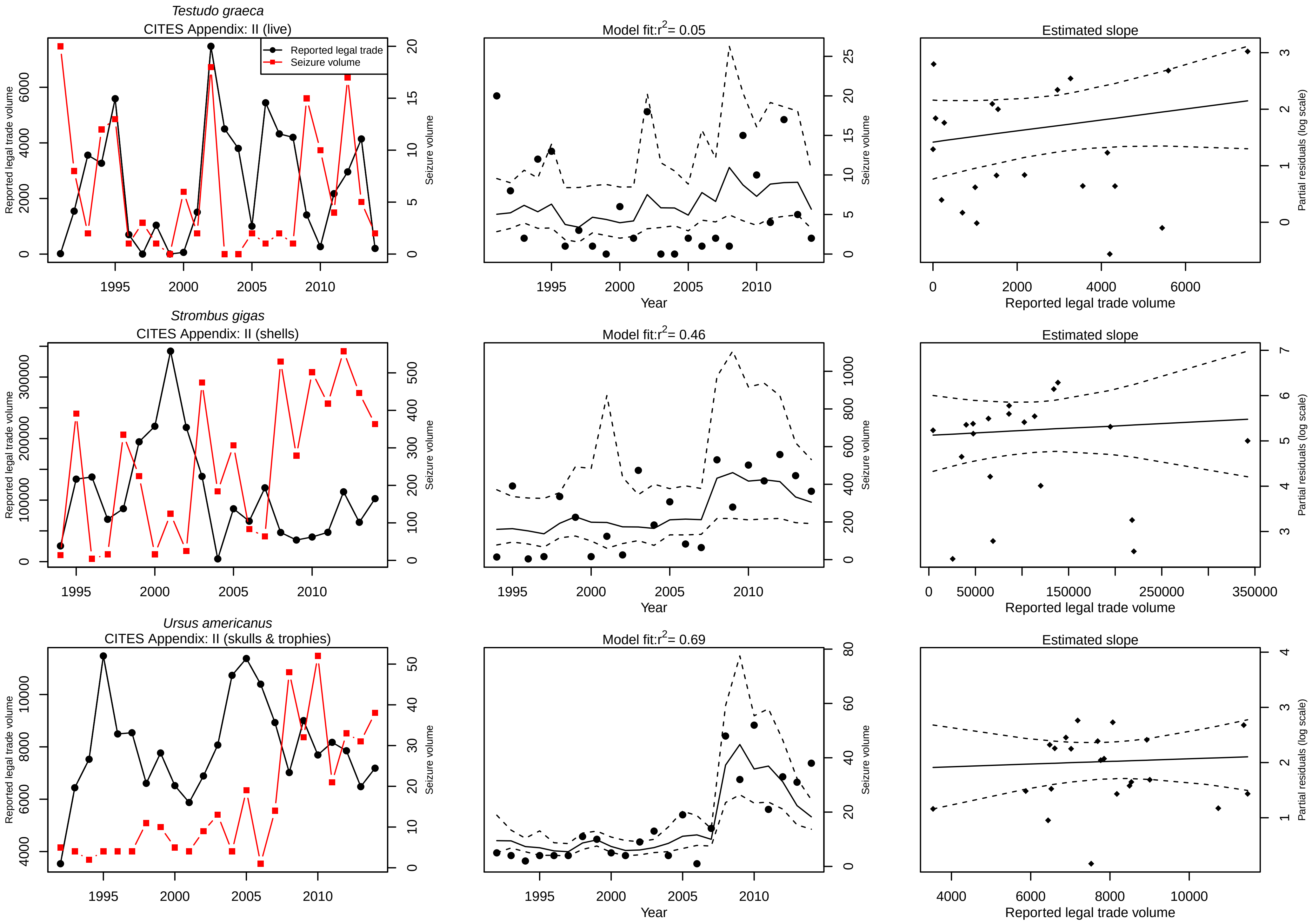

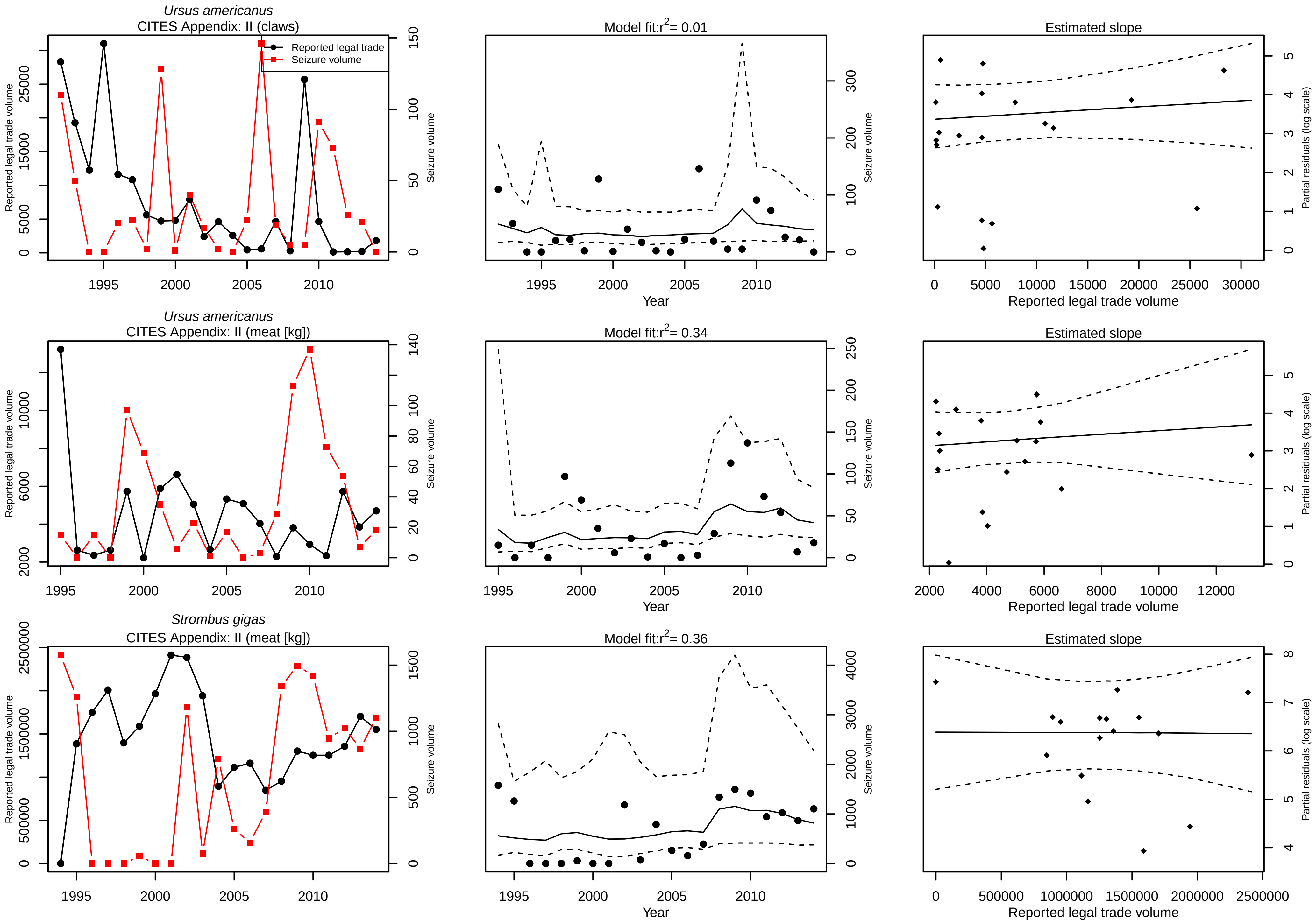

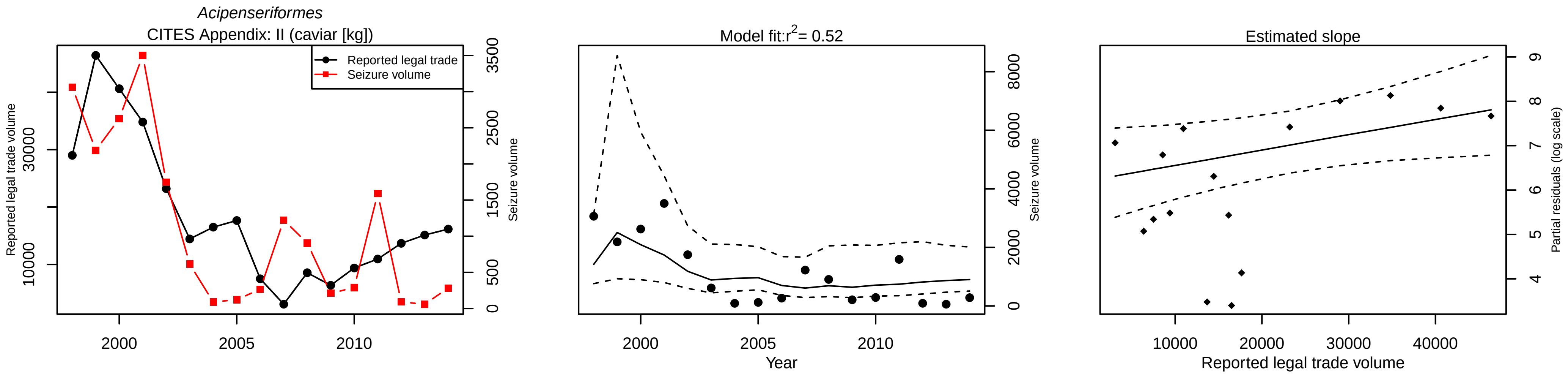

### Figure S2

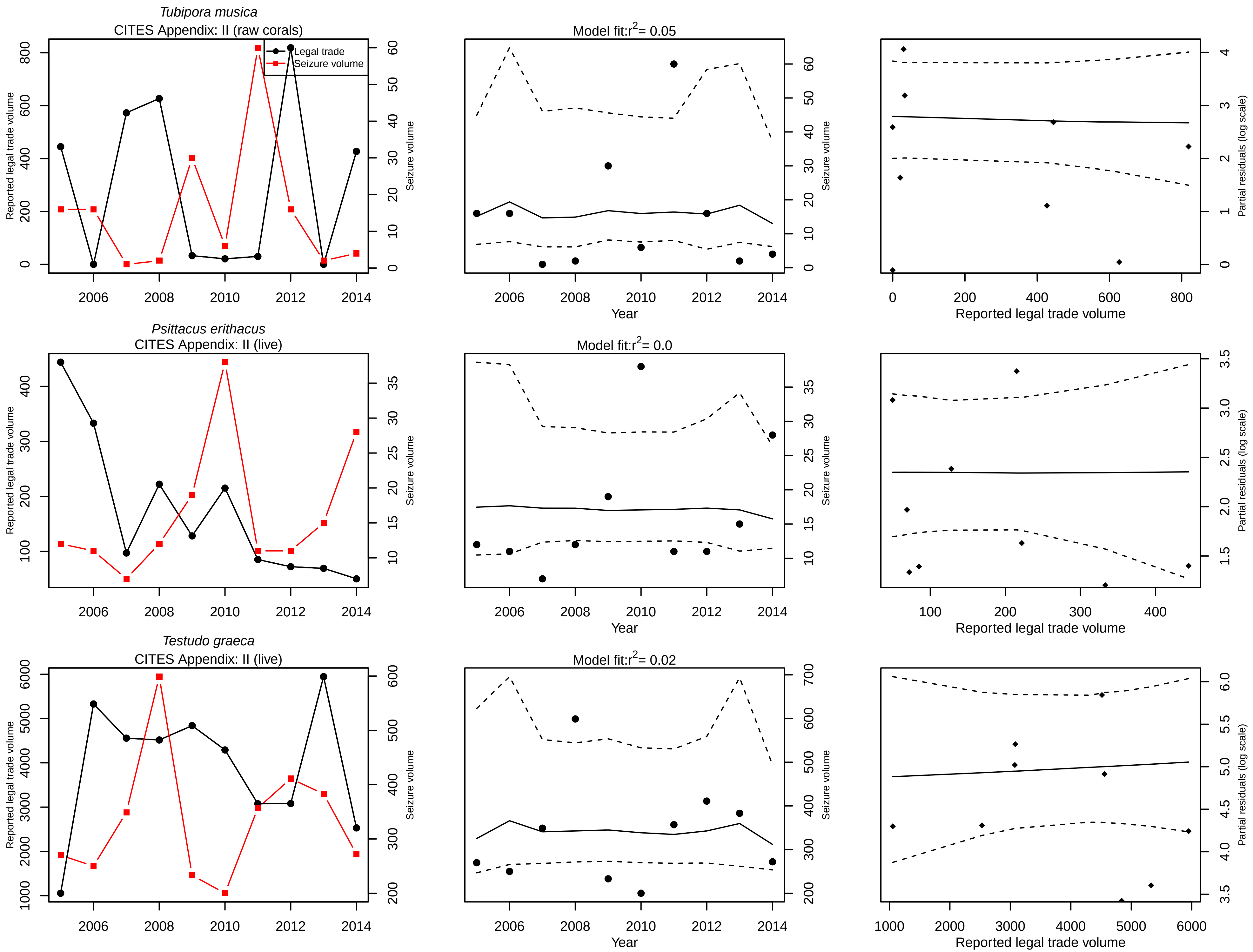

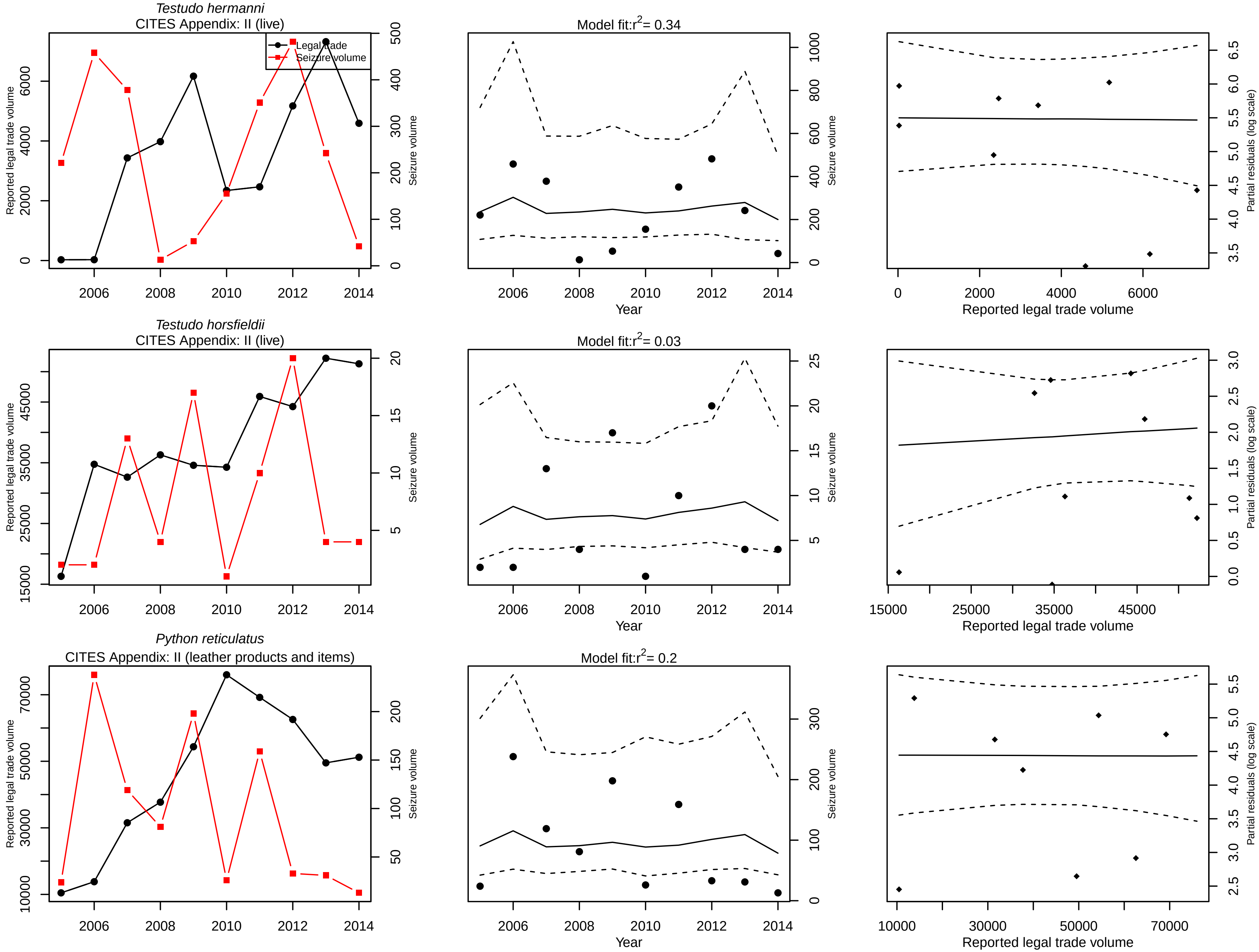

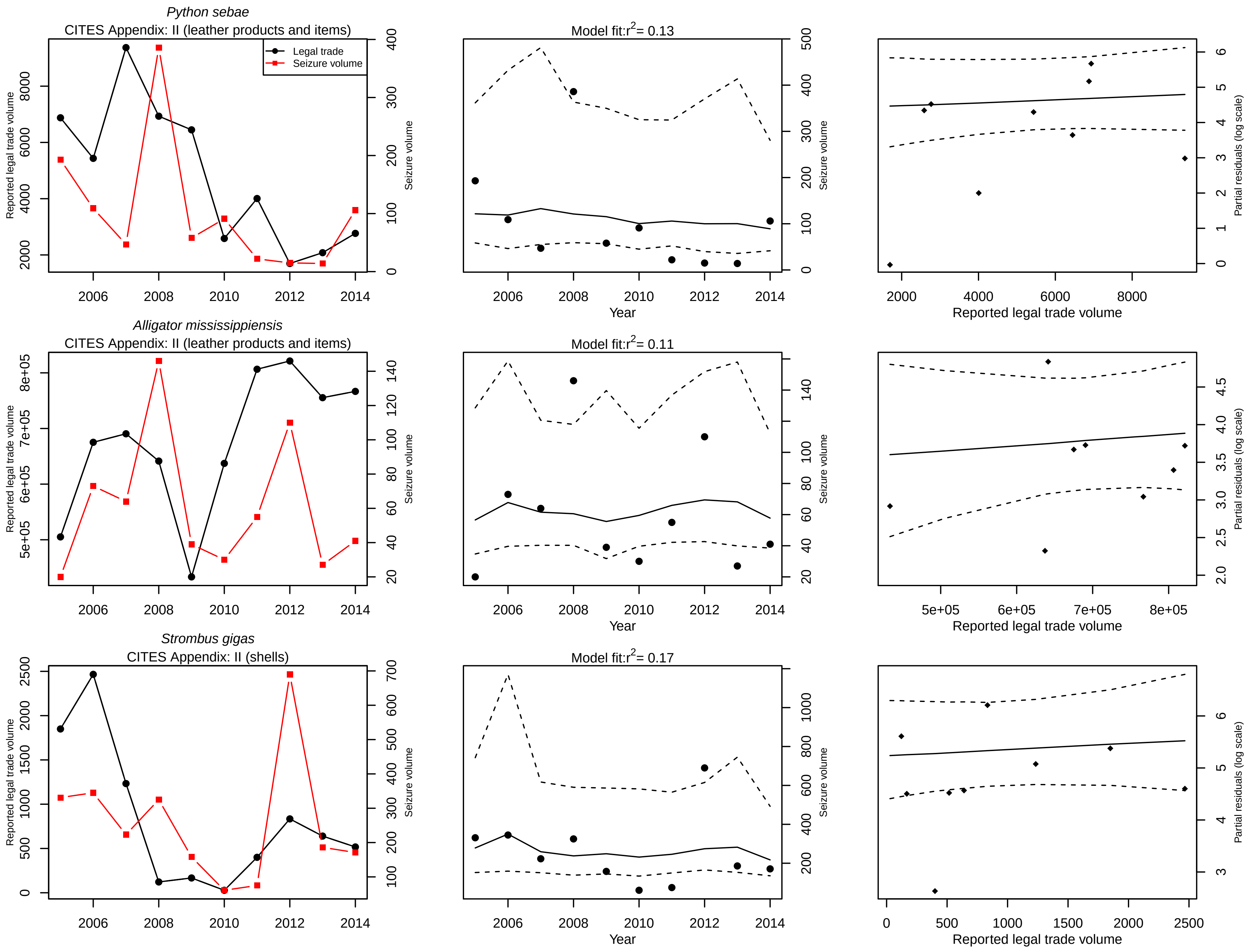

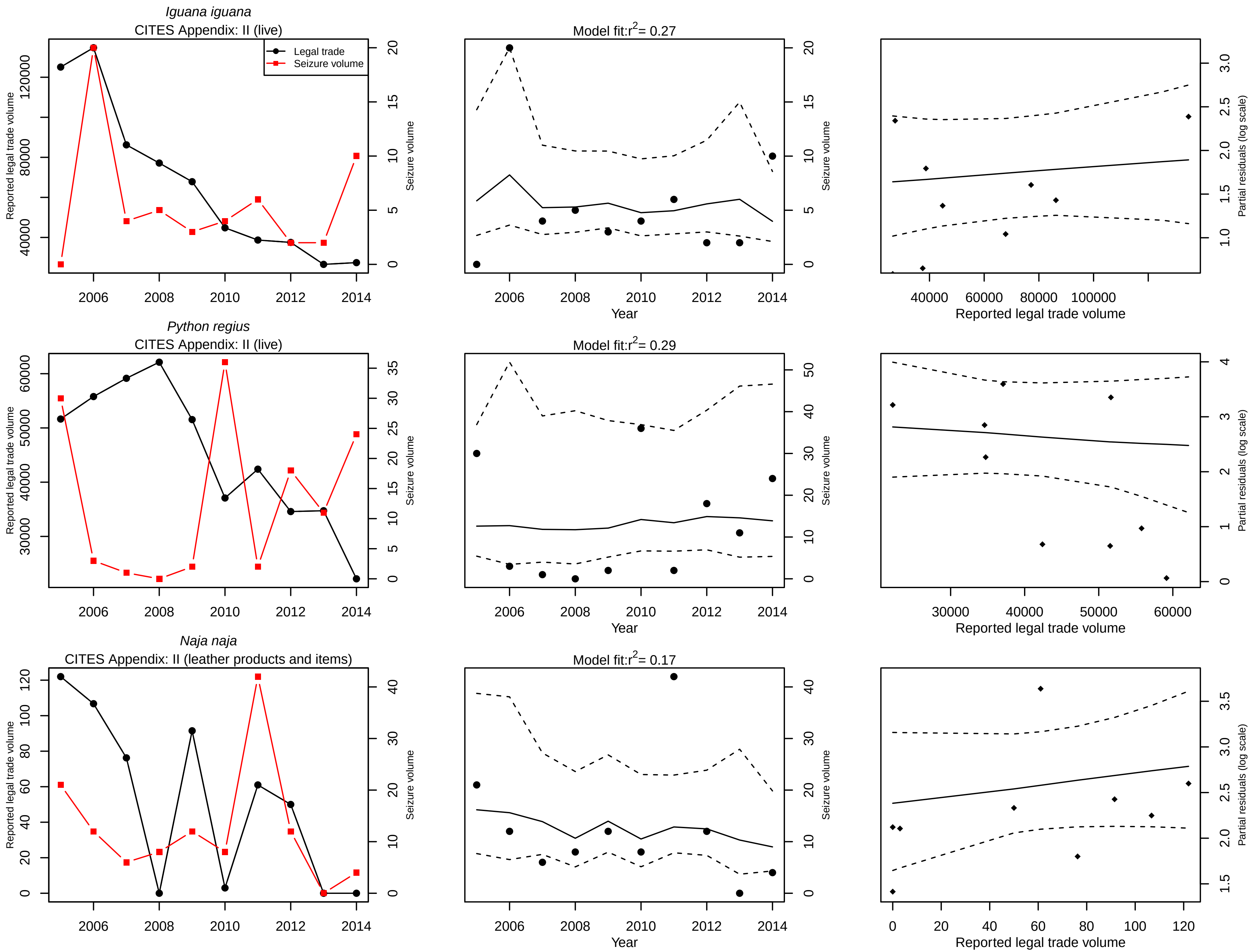

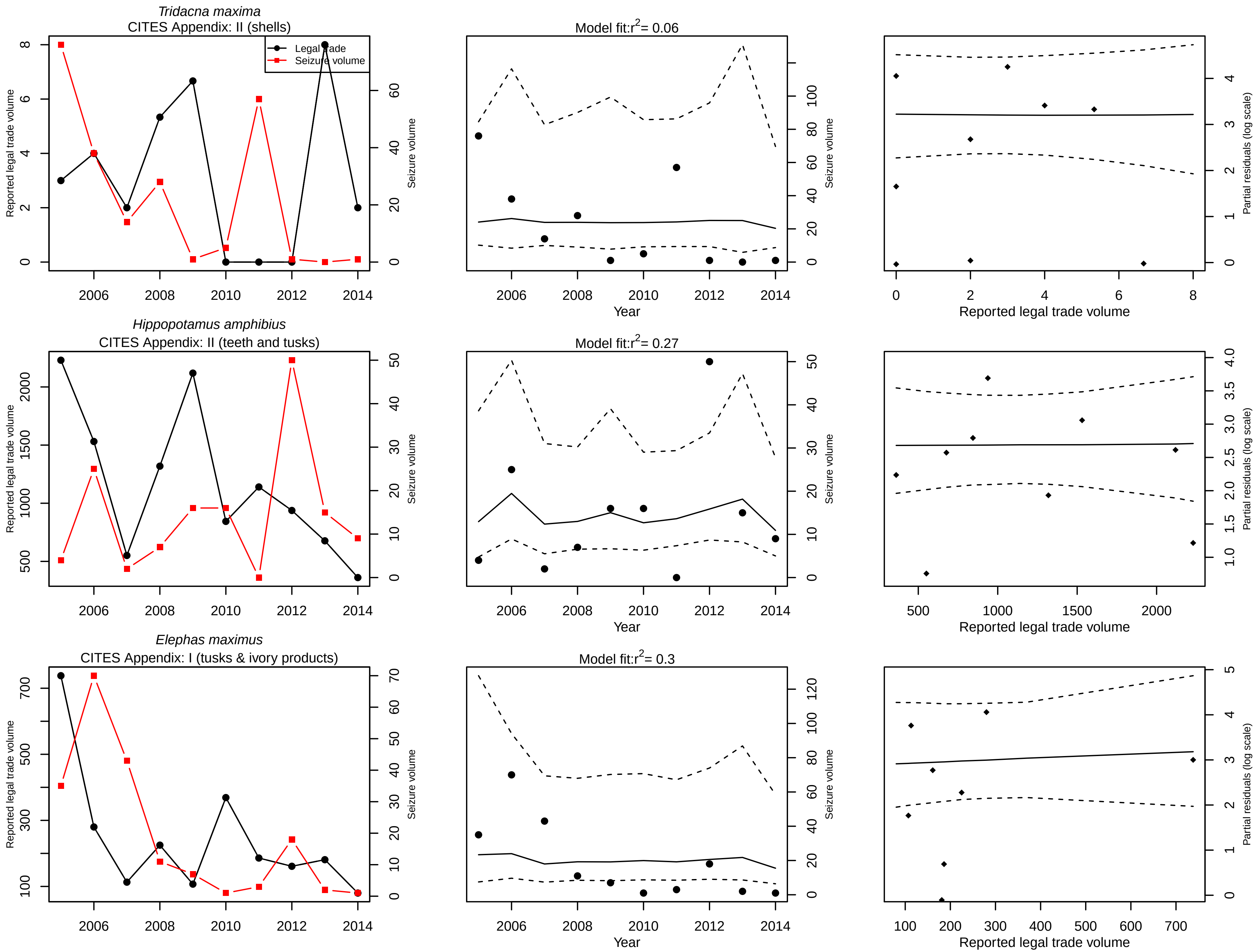

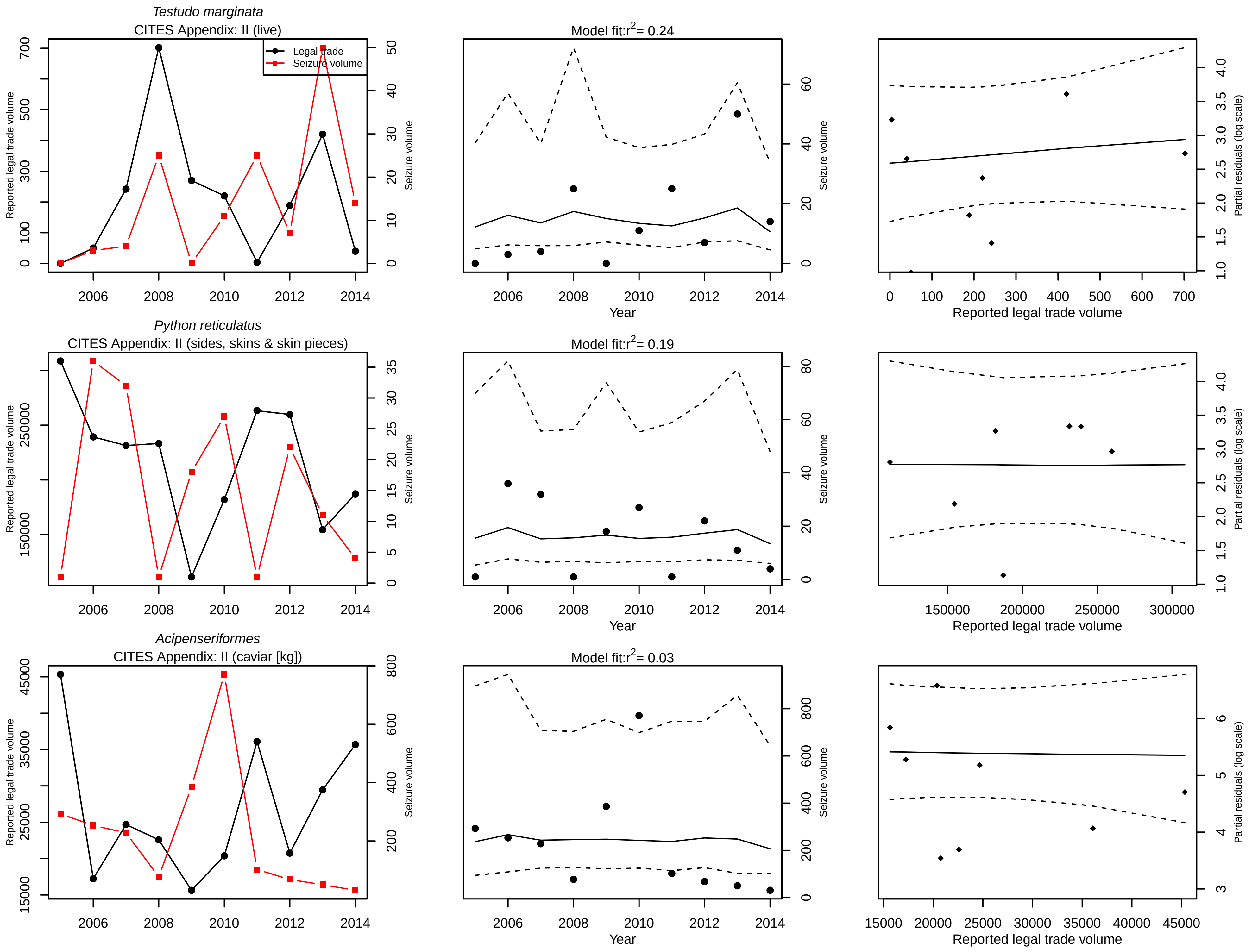

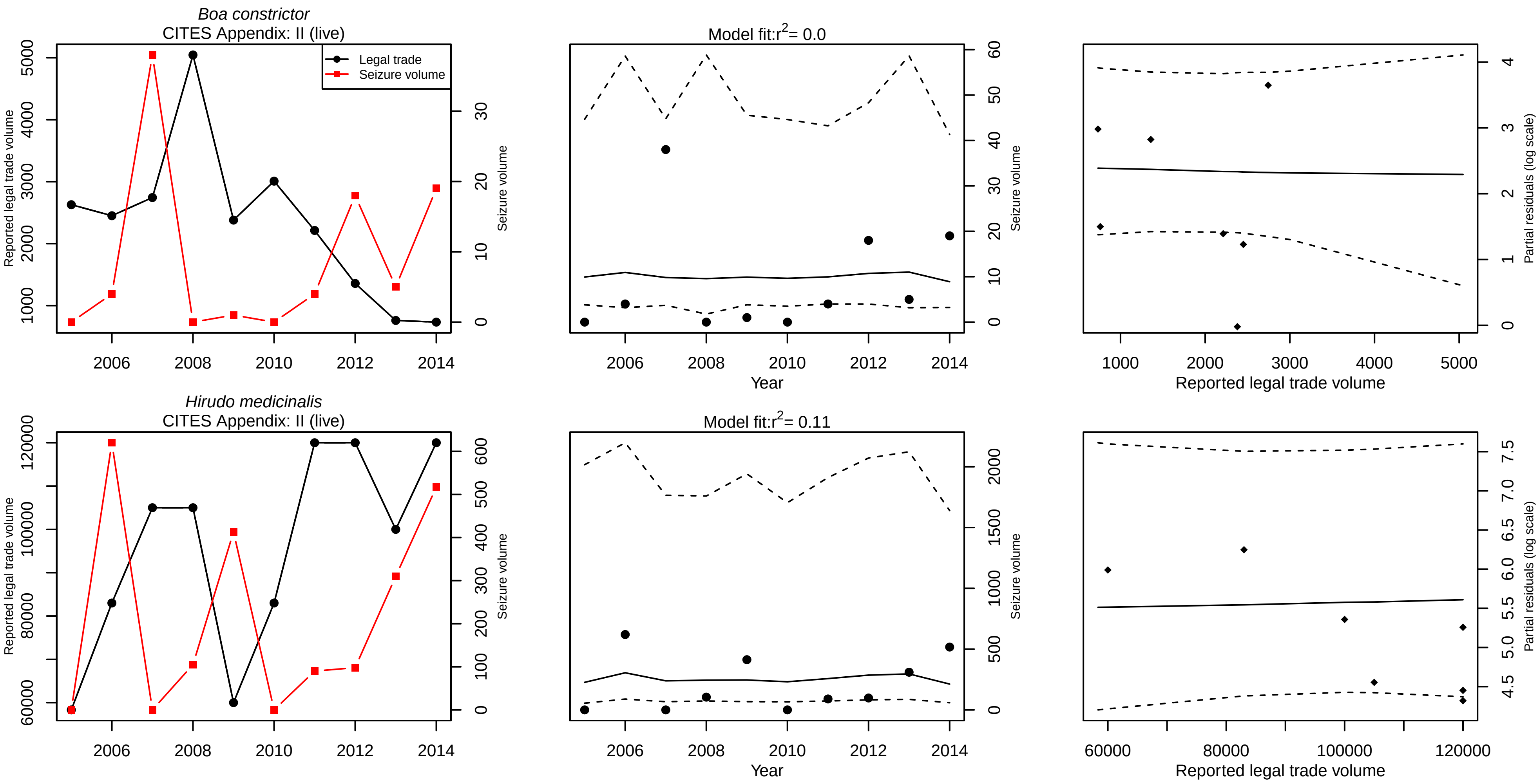
